# Coordinated crosstalk between microtubules and actin by a spectraplakin regulates lumen formation and branching

**DOI:** 10.1101/2020.05.09.085753

**Authors:** Delia Ricolo, Sofia J. Araújo

**Affiliations:** Department of Genetics, Microbiology and Statistics, School of Biology, University of Barcelona, 08028 Barcelona, Spain; Institute of Biomedicine University of Barcelona (IBUB), Barcelona, Spain

**Author notes:** Corresponding author. email: S. J. Araújo.

**Keywords:** lumen, subcellular, branching, Drosophila, trachea, short-stop, shot, tau

## Abstract

The establishment of branched structures by single cells involves complex cytoskeletal remodelling events. In *Drosophila*, epithelial tracheal system terminal cells (TCs) and dendritic arborisation neurons are models for these subcellular branching processes. During tracheal embryonic development, the generation of subcellular branches is characterized by extensive remodelling of the microtubule (MT) network and actin cytoskeleton, followed by vesicular transport and membrane dynamics. We have previously shown that centrosomes are key players in the initiation of subcellular lumen formation where they act as microtubule organizing centres (MTOCs). However, not much is known on the events that lead to the growth of these subcellular luminal branches or what makes them progress through a particular trajectory within the cytoplasm of the TC. Here, we have identified that the spectraplakin *Short-stop* (*Shot*) promotes the crosstalk between MTs and actin, which leads to the extension and guidance of the subcellular lumen within the TC cytoplasm. Shot is enriched in cells undergoing the initial steps of subcellular branching as a direct response to FGF signalling. An excess of Shot induces ectopic acentrosomal branching points in the embryonic and larval tracheal TC leading to cells with extra subcellular lumina. These data provide the first evidence for a role for spectraplakins in subcellular lumen formation and branching.

## INTRODUCTION

Cell shape is intrinsically connected with cell function and varies tremendously throughout nature. Tissue and organ morphogenesis rely on cellular branching mechanisms that can be multicellular or arise within a single-cell. Through extensive cellular remodelling, this so-called single-cell or subcellular branching, transforms an initially relatively symmetrical unbranched cell into an elaborate branched structure. These cellular remodelling events are triggered by widespread cytoskeletal changes and cell membrane growth, which allow these branched cells to span very large areas and accomplish their final function. Despite this clear link between morphology and function, not much is known about the signalling events that trigger the formation of these subcellular branches or what makes them choose a particular trajectory within the cytoplasm of the cell.

In *Drosophila melanogaster*, tracheal system terminal cells (TCs) and nervous system dendrites are models for these subcellular branching processes. During tracheal embryonic through larval development, the generation of single-cell branched structures by TCs is characterized by extensive remodelling of the MT network and actin cytoskeleton, followed by vesicular transport and membrane dynamics (1–3). During embryonic development, TCs, as tip-cells, lead multicellular branch migration and extension in response to Bnl-Btl signalling, which induces the expression of *Drosophila* Serum Response Factor (DSRF/*blistered (bs)*) and its downstream effectors (4, 5). Although epithelial in origin, TCs do not have a canonical apical-basal polarity, and, as they migrate, extend numerous filopodia on their basolateral membrane, generating transient protrusive branches at the leading edge (6). As a consequence, they display a polarity similar to that of a migrating mesenchymal cell (7).

While migrating and elongating, the TC invaginates a subcellular tube from its apical membrane, at the contact site with the stalk cell (1). The generation of this *de novo* subcellular lumen can be considered the beginning of the single-cell branching morphogenesis of this cell, which continues throughout larval stages to generate an elaborate single-cell branched structure with many subcellular lumina (3).

We have previously shown that centrosomes are key players in the initiation of subcellular branching events during embryogenesis. Here, they act as microtubule organizing centres (MTOCs) mediating the formation of single or multiple branched structures depending on their numbers in the TC (8). Centrosomes organise the growth of MT-bundles towards the elongating basolateral edge of the TC. These MTs have been suggested to serve both as trafficking mediators, guiding vesicles for delivery of membrane material, and as mechanical and structural stabilizers for the new subcellular lumen (3). Actin filaments are present at the growing tip, the basolateral and the luminal membrane of the TC, and actin-regulating factors such as DSRF, Enabled (Ena) and Moesin (Moe) have been shown to contribute to TC morphogenesis (1, 9, 10). During TC elongation, the lumen extends along with the cell, stabilizing the elongating cell body and maintaining a more or less constant distance between its own tip and the migrating tip of the cell (1). At the TC basolateral side, a dynamic actin pool integrates the filopodia and aligns the growing subcellular tube with the elongation axis (11–13). Together, MT-bundles and the basolateral actin pool are necessary for subcellular lumen formation (1). However, not much is known on how these two cytoskeletal structures are coordinated within the TC.

By the time the larva hatches, TCs have elongated and grown a full-length lumen, which becomes gas-filled along with the rest of the tracheal system. In the larva, terminal cells ramify extensively and form many new cytoplasmatic extensions each with a membrane-bound lumen creating tiny subcellular tubes that supply the targets with oxygen (14–16). At larval stages, sprouting and extension of new branches in response to local hypoxia is generally considered to occur by essentially the same molecular mechanisms as the initial tube invagination and cell extension in the embryo (2, 17). However, not much is known about how hypoxic signalling is transduced into cytoskeletal modulation to achieve the single-cell branching morphogenesis of the TC. Also, what coordinates the crosstalk between microtubules and actin at the basolateral growing tip, how cell elongation is stabilized by lumen formation and how both processes remain coordinated is still poorly understood in both embryonic and larval TCs.

Spectraplakins are giant conserved cytoskeletal proteins with a complex multidomain architecture capable of binding MTs and actin. They have been reported to crosslink MT minus-ends to actin-networks, making MT-bundles more stable and resistant to catastrophe (18). Loss of spectraplakins has been shown *in vivo* to have remarkable effects on microtubule organization, cell polarity, cell morphology, and cell adhesion (19, 20). *Drosophila* has a single spectraplakin, encoded by *short-stop* (*shot*) (20–22). *shot* mutants display pleiotropic phenotypes in wing adhesion, axon and dendrite outgrowth, tracheal fusion, muscle-tendon junction, dorsal (23)closure, oocyte specification and patterning, photoreceptor polarity and perinuclear microtubule network formation (21, 24–29). Shot has been shown to bind both the microtubule plus-end-binding EB1 and the microtubule minus-end-binding protein Patronin, required for the establishment of acentrosomal microtubule networks (26, 27, 30). It also has been shown to bind actin and to crosslink MTs and actin contributing to cytoskeletal organization (24).

In the present study, we uncover a novel role for the spectraplakin Shot in subcellular lumen formation and branching. Our results show that *shot* loss-of-function (LOF) leads to cells deficient in *de novo* subcellular lumen formation at embryonic stages. We show that Shot promotes the crosstalk between microtubules and actin, which leads to the extension and guidance of the subcellular lumen within the TC cytoplasm. We observe that Shot levels are enriched in cells undergoing the initial steps of subcellular branching as a direct response to FGF signalling. And an excess of Shot induces ectopic acentrosomal branching points in the embryonic and larval tracheal TC leading to cells with extra subcellular lumina. Furthermore, we find that Tau protein can functionally replace Shot in subcellular lumen formation and branching.

## RESULTS

### Loss of Shot causes defects in *de novo* subcellular lumen formation

Shot is expressed during *Drosophila* development in several tissues such as the epidermis, the midgut primordia, the trachea and the nervous system (25, 31). We began by analysing the effect of *shot* LOF during TC subcellular lumen formation. To do so, we analysed dorsal (DB) and ganglionic branch (GB) TCs at late stages of embryogenesis (st. 16) (Fig. 1 A, B).

**Figure 1.**
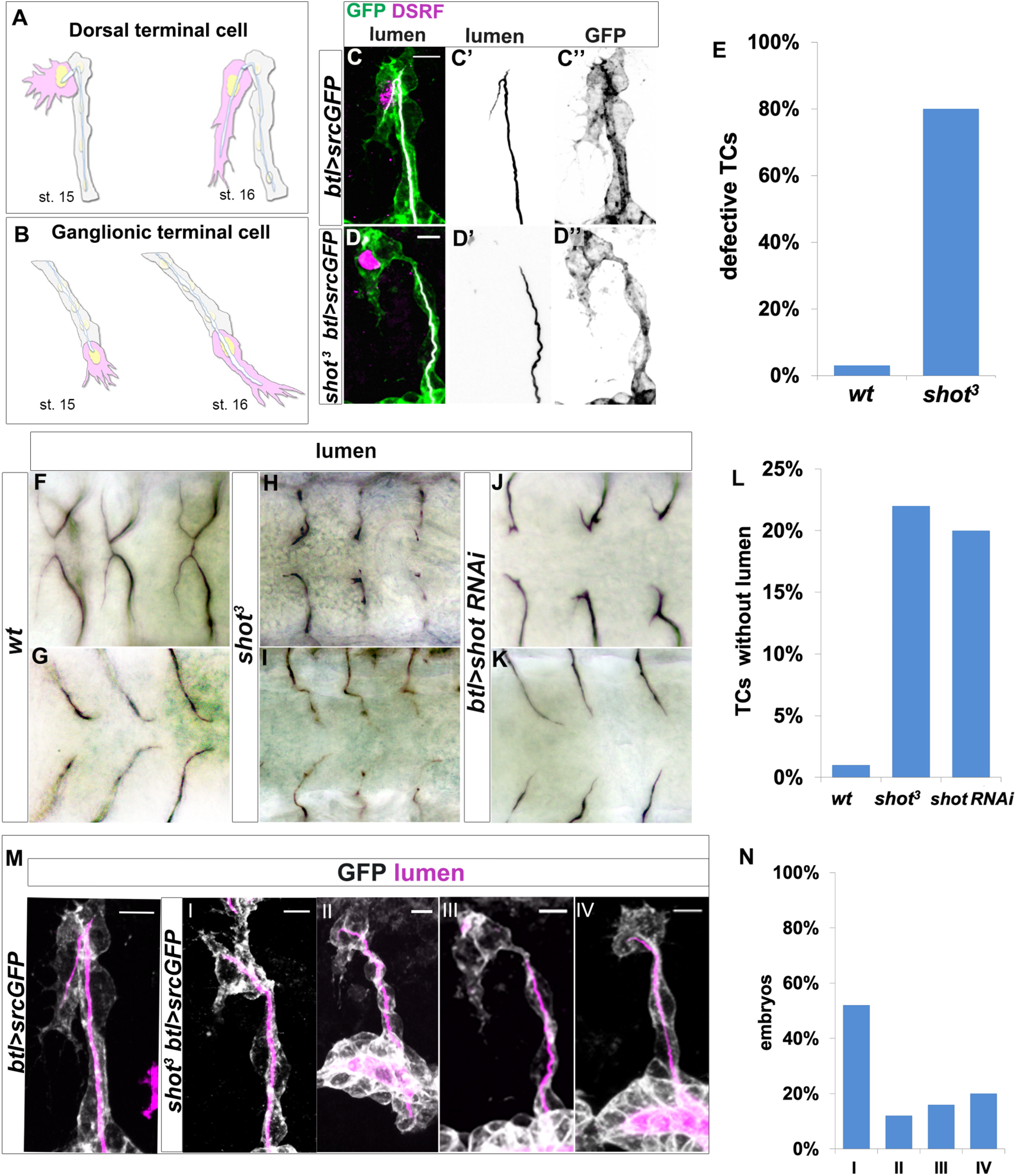
*shot* loss of function induces defects in subcellular lumen formation. (A-B) Representation of dorsal and ganglionic TCs from st. 15 to st. 16 (DB and GB in grey, TC in pink). At st.15, the TC (cytoplasm in pink, nucleus in yellow, basal membrane in grey, apical membrane in blue and lumen in white) emits filopodia in the direction of cell migration and elongates; apical membrane grows in the same direction giving rise to the outline of the subcellular lumen. As it extends, the subcellular lumen is filled of chitin (white). At the end of st.16 the TC has elongated and the subcellular lumen is formed inside the cytoplasm, creating a new apical surface in the TC. (C-D) DBs at st.15 of *btl>srcGFP* (control, C) and *shot^3^; btl>srcGFP* (D) fixed embryos stained with GFP to visualize tracheal cells, green in C and D, grey in C’’ and D’’, CBP to visualize the lumen, white in C and D, black in C’ and D’ and DSRF in magenta. Anterior side is on the left and dorsal is up, scale bars 5 μm. (E) Quantification of defective TCs in *shot^3^* and *wt* (n= 20 embryos, 400TCs). (F-K) DBs (F, H, J dorsal view) and GBs (G, I, K ventral view) of fixed embryos stained with anti-Gasp antibody at st.16 of *wt* (F and G), *shot^3^* (H and I) and *btl>shotRNAi* (J and K) (L) Quantification of TCs (genotype indicted) without subcellular lumen (*wt* n=400, *shot^3^*=400*, btl>shotRNAi* n=300). (M-N) Different types of TC mutant phenotypes were produced in *shot* LOF conditions. (M) Dorsal branches of *btl>srcGFP* control (*wt*) and *shot^3^* embryos stained with GFP (grey) to visualize membrane and CBP (in magenta) to visualize the lumen. Anterior side is on the left and dorsal side is up. Scale bars 5 μm. (I) TC partially elongated with formed lumen but with wrong directionality (52%); (II) the elongation was stopped prematurely and a primordium of subcellular lumen was formed (12%); (III) the cell elongated partially but the lumen was completely absent (16%); and (IV) the cell was not able to elongate and the lumen was completely absent (20%). (N) Quantification of the different types of TC mutant phenotypes reported as I-IV (n=25 TCs).

The *shot^3^* null mutant TC phenotype consisted in subcellular lumen elongation defects with a penetrance of 80% (Fig. 1 C, D and F-I and E). This phenotype resembled the previously reported for *blistered* (*bs*) mutants (9). *bs* encodes the transcription factor DSRF that regulates TC fate induction in response to Bnl-Btl signalling (9, 32). However, we observed that DSRF was properly accumulated in *shot^3^* TC nuclei (Fig. 1 D), discarding a possible effect of Shot in TC fate induction.

To analyse if the *shot* phenotype was cell autonomous, we expressed *shot* RNAi to knock-down Shot in all tracheal cells and found that, like in null mutant conditions, 80% of TCs analysed (n=300) at the tip of the dorsal branches (n=150) or ganglionic branches (n=150) were affected in subcellular lumen formation (Fig. 1 J, K). Of these, 20% did not develop a terminal lumen at all (Fig. 1 L).

*shot^3^* embryonic TC lumen phenotypes range in expressivity from complete absence of subcellular lumen to different lengths of shorter lumina (Fig. 1 M, N). When quantified, out of the 80% of embryos that showed a TC luminal phenotype, 36% of TCs did not elongate a subcellular lumen at all and 64% failed to accomplish a full-length lumen (300 ganglionic TCs and 300 dorsal TCs) (Fig. 1 M, N). Taken together, these results indicated that Shot is involved in *de novo* subcellular lumen formation and elongation.

### Shot overexpression induces extra subcellular branching independently of the centrosome

Having observed that Shot was necessary for subcellular lumen formation and extension, we hypothesised that Shot overexpression (ShotOE) would induce extra subcellular branching events. Indeed, analysis of full-length ShotOE (*shotA-GFP*) in tracheal cells revealed that increasing Shot concentrations induced Extra-Subcellular Lumina (ESL) in GB and DB TCs (Fig. 2 A-C, G, H). Since MTs and actin are essential for subcellular lumen formation (1), we then asked whether supernumerary luminal branching was due to the MT- or the actin-binding domains present in the Shot molecule (33, 34). To this end, we overexpressed an isoform of Shot (ShotC-GFP) with a deletion of the first calponin domain (Fig. 2 M, N), resulting in a shorter acting binding domain (ABD), which binds actin very weakly or not at all (22, 25). The tracheal overexpression of *shotA* induced phenotypes in 95% of the embryos (n=20), with an average of 2 TC bifurcations *per* embryo (n=400). *shotC* overexpression induced phenotypes in 90% of the embryos (n=20), with an average of 2 TC bifurcations *per* embryo (n=400) (Fig. 2 G). In all cases, we could detect more MT bundles in TCs, associated with the ESLs (Figure S1). ShotA-GFP and ShotC-GFP displayed different localizations within the TC. Full-length ShotA-GFP localization can be detected at the cell-junctions, around the crescent lumen, in structures resembling MT-bundles, and throughout the cytoplasm, whereas ShotC-GFP localized more to the MT/lumen region, together with MT-bundles (Fig. S1), in agreement with the lack of actin-binding capability of ShotC isoform. Interestingly, we observed a highly ramified subcellular lumen when higher amounts of ShotC were expressed in tracheal cells (Fig. 2 J) suggesting that the effect of ShotOE in subcellular lumen branching was dosage dependent.

**Figure 2.**
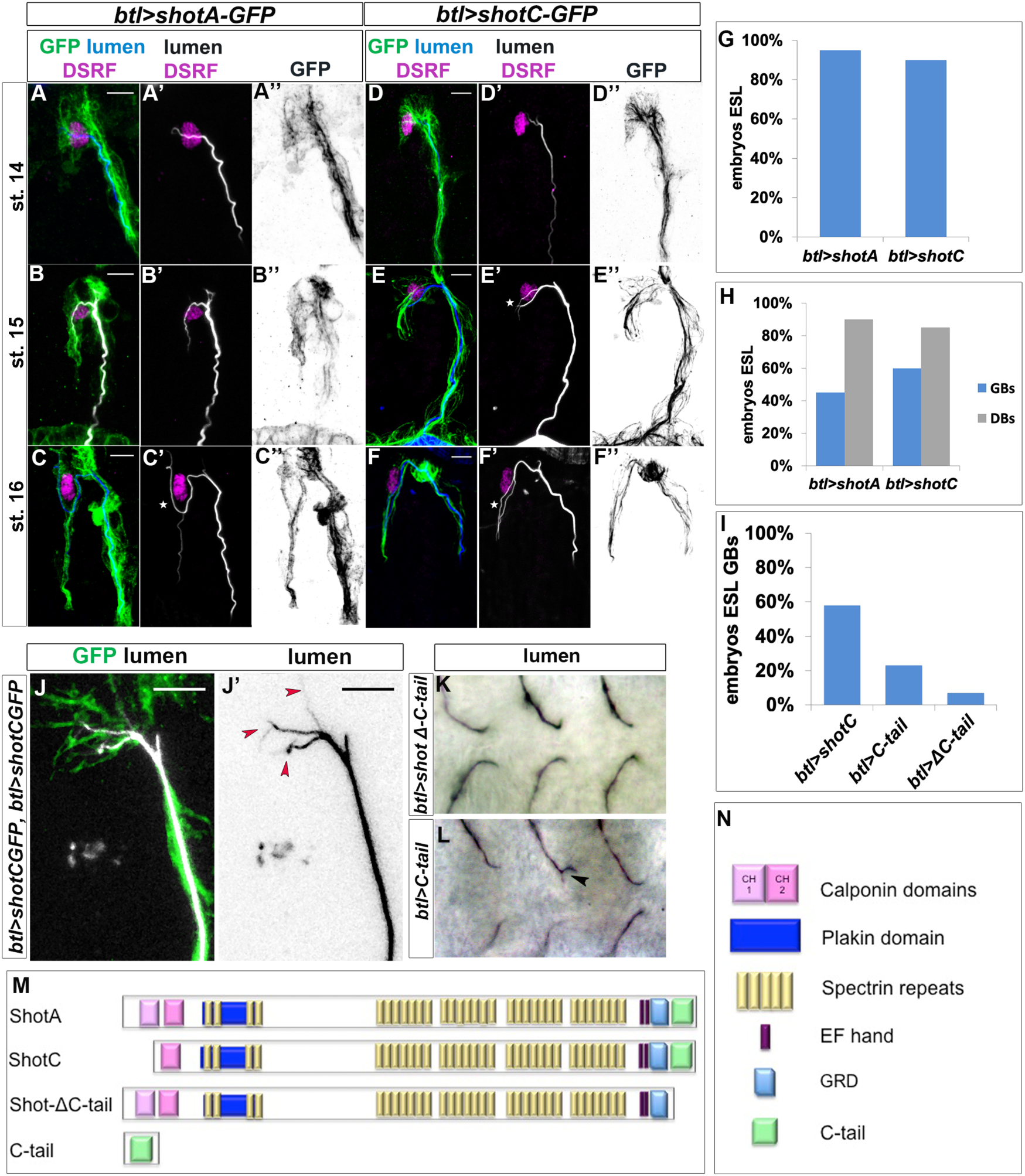
*ShotOE* induces luminal branching through its microtubule binding domain. Lateral view of DB tip cells from st.14 to st.16, of *btl>shotA-GFP* embryos (A-C) and *btl>shotCGFP* (D-F). Embryos were stained with GFP (green in A-F and grey A’’-F’’) to visualize Shot-GFP, DSRF to mark the TC nuclei (in magenta) and CBP to stain the chitinous lumen (blue in A-F and white A’-F’). Both overexpressing conditions induced ESLs (white stars). Note that GFP was more distributed throughout the TC cytoplasm of embryos overexpressing *shotA*, and more organized in bundles in TCs overexpressing *shotC*. Anterior side of embryo is on the left and dorsal side up. Scale bars 5μm. (G) Penetrance of ESL phenotype of embryos overexpressing *shotA* (n=20 embryos, 400 TCs) and *shotC* (n=20 embryos, 400 TCs) in all trachea cells, displaying at least one TC affected, considering both GBs and DBs. (H) Distribution of ESL phenotype in GBs (n=200 TCs) blue column and DBs (n = 200 TCs) grey column. (J) ESL phenotype induced by Shot is dosage dependent. Example of dorsal TC of an embryo overexpressing two copies of *btl>shotC-GFP*, stained with anti-GFP and CBP. Red arrows indicate extra subcellular lumen branching. Note that the ESL are very thin and follow Shot positive bundles detected with GFP. Anterior side is on the left, dorsal midline is on the top. Scale bars 5μm. (I, K, L) The C-tail domain is involved in ESL formation. (I) Percentage of embryos overexpressing *shotC*, *shotΔCtail and C-tail* in the tracheal system displaying GB ESLs (n= 40 embryos, 400TCs each genotype). Tips of GB TCs from *btl>shotΔCtail* embryos with a single subcellular lumen each (K) and *btl>C-tail* (L) in which one TC is bifurcated; stained with anti-Gasp (ventral view, anterior side of the embryo is on the left). (M) Schematic representation of different Shot constructs used and (N) Spectraplakin protein domains.

Tracheal overexpression of *shotC* phenocopied that of *shotA* in inducing ESLs (Fig. 2 D-F, G, H), suggesting that the ABD is not necessary for the induction of additional luminal branching events. In order to clarify this, we used two other isoforms of Shot: *shot∆Ctail*, lacking the C-terminal MT-binding domain, and *shotCtail*, a truncated form containing only the C-terminal MT-binding domain (35) (Fig.2 M, N). Whereas overexpressing *shotΔ-Ctail* in TCs we could only detect a branching phenotype in 7% of embryos analysed (n=40), (Fig. 2 K), overexpression the C-tail domain alone induced TCs with extra branching in 23% of the embryos (n=40) (Fig. 2 L), indicating that the C-tail alone was sufficient to induce ESLs in TCs. Taken together these results using different Shot isoforms, lead us to conclude that the Shot MT-binding domain alone is sufficient for the extra branching events observed in ShotOE TCs.

ESLs were previously observed when higher numbers of centrosomes were present in TCs (8). We therefore asked if the observed extra branching phenotypes could be due to supernumerary centrosomes induced by ShotOE in TCs. Consequently, we quantified the number of centrosomes in the TCs of ShotOE embryos. In control TCs we detected an average of 2,3 ± 0,5 (n=33) centrosomes per TC, and in ShotOE 2,2 ± 0,2 centrosomes per TC (n=33) (Fig. 3 A, B, D). In both conditions, and as previously described (8), this centrosome-pair was detected at the apical side of the TCs (Fig. 3 A, B). Besides, analysing ShotOE TCs at embryonic st.15, (n=16) we could detect that the ESL arose from the pre-existing subcellular lumen, distally from the centrosome pair (Fig. 3 B’ arrow). These data indicate that ShotOE did not change TC centrosome-number and induced ESL by a distinct mechanism from centrosome duplication.

**Figure 3.**
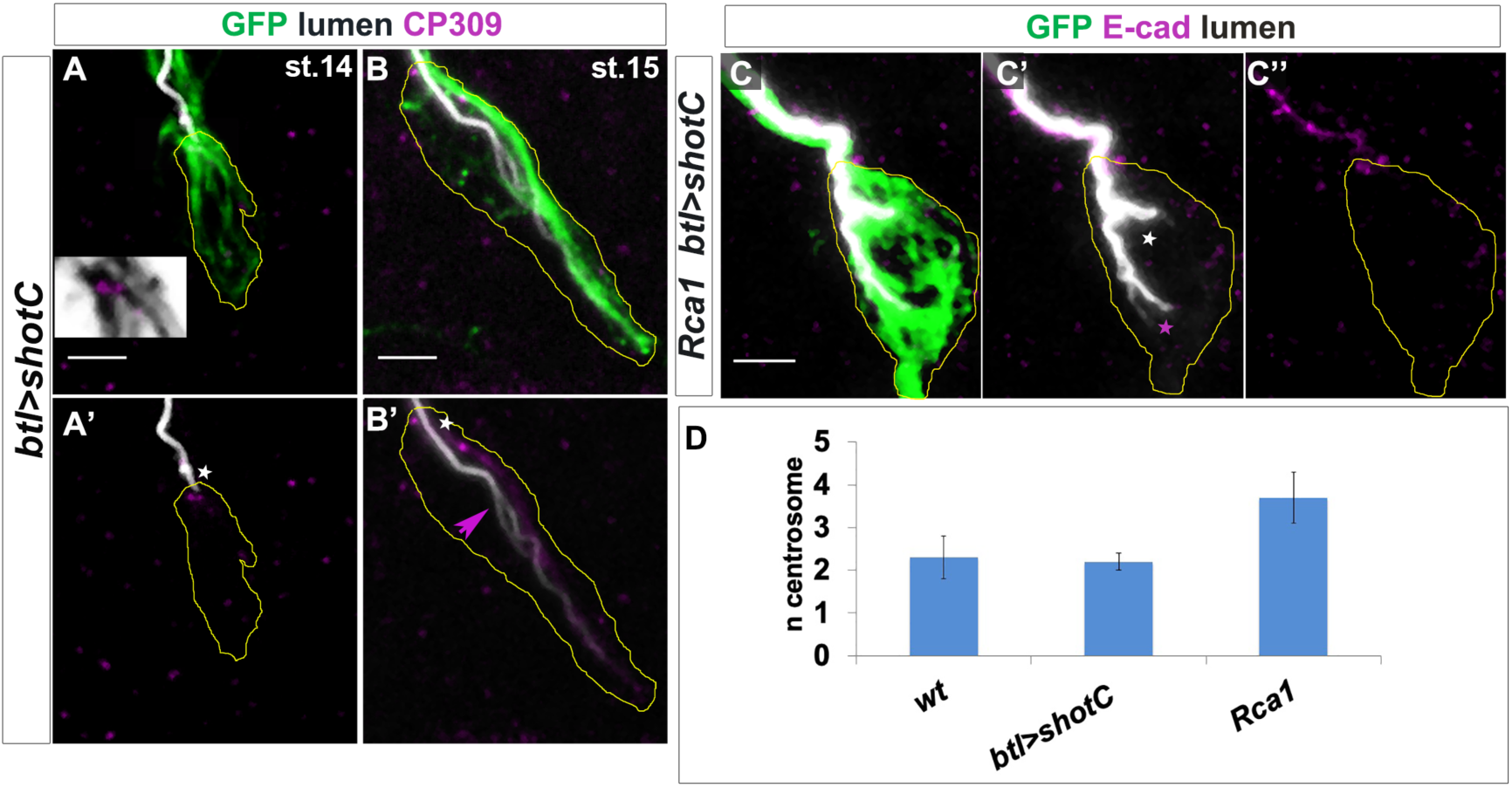
ESL induction by *ShotOE* is not associated with centrosome amplification. (A, B) GB TC of st. 14 (A) and st. 15 (B) *btl>shotC-GFP* embryos stained with CBP to mark lumen (white), GFP to visualize Shot (green) and CP309 to mark centrosomes (magenta); the outline of TCs is drawn in yellow. The box in A is a digital magnification showing the TC centrosome pair (magenta) and GFP positive Shot bundles (in grey) emanating from centrosomes. White stars indicate apically localized centrosomes. In B’ the subcellular lumen (magenta arrow) is bifurcated far from the centrosome-pair, from the pre-existing lumen. (C) GB tips from *Rca1; btl>shotC-GFP* embryos at st. 15, stained with CBP (in white) to visualize the lumen and E-cadherin (in magenta) to recognize the apical junction. Anterior side of the embryo is on the left and ventral is down. Scale bar 2 μm. In these cases, two types of luminal bifurcations are detected: one from the apical junction (white asterisk), caused by *Rca1* supernumerary centrosomes and another arising from the pre-existing lumen (magenta asterisk), caused by ShotOE. (D) Quantification of centrosome number in *wt, btl>shotC* and *Rca1* embryos ± SEM.

In contrast with ShotOE alone (Fig. 3 B’), in *Rca1* mutants the bifurcated subcellular lumen arose from the apical junction and continued to extend during TC development (8). When we analysed the luminal origins in *Rca1*, ShotOE conditions, both types of ESL where detected in the same TC in 25% of the cases (n=12). In the same TCs two types of ESL were generated, one from the apical junction and another sprouted from the pre-existing lumen distally from the junction (Fig. 3 C, asterisks). In addition, the effect of *Rca1* LOF and ShotOE was additive in producing TCs with a multiple-branched subcellular lumen (Fig. S2). These morphological ESL differences suggested that *Rca1* and *shot* operate in different ways in the *de novo* formation and branching of the subcellular lumen.

### Shot interacts with stable microtubules and actin

Spectraplakin expression is critical in cells that require extensive and dynamic cytoskeleton reorganization, such as epithelial, neural, and migrating cells. Loss of spectraplakin function leads to a variety of cellular defects due to disorganised cytoskeletal networks (36). In a plethora of tissues and in cultured S2 cells, Shot can physically interact with different cytoskeletal components (23, 24, 37). Therefore, we investigated Shot localization and its interaction with MTs and actin in *wt* TCs.

We analysed live embryos using time-lapse imaging, and observed that Shot localization was extremely dynamic throughout subcellular lumen formation. We could detect Shot in the apical TC junction as well as extending together with the growing subcellular lumen (Movie 1 and Fig. S3 A). It was apparent that Shot localized dynamically with the growing luminal structures, showing a strong localization at the middle/tip of the extending TC (Movie 1 and Fig. S3 A).

In *wt* conditions F-actin and the actin-binding protein Moe concentrate strongly at the tip of the TC, but are also detected in the TC cytoplasm, and these different actin populations have been shown to be important for subcellular lumen formation and extension (1, 11). During TC elongation, MTs polymerise from the centrosome pair at the apical junction towards the tip of the cell, reaching the area of high Moe and F-actin accumulation (1, 8). So, we next analysed Shot localization in relation to the dynamically localized actin core present in the cytoplasm and at the tip of the migrating TC in live embryos (Movie 2, 3). In both GB and DB TCs we could detect a dynamic interaction between Shot and Moe (Movie 2 and Fig. S3 B) and Actin (Movie 3 and Fig. S3 C). As the lumen extended, Shot interacted with different actin populations, namely the actin core and basal, filopodial actin (Fig. S3 B, C).

We followed these analyses, observing endogenous Shot in fixed and antibody stained embryos. At early stages, when TCs started to elongate, we detected Shot co-localizing with Moe at the tip of the TC (Fig. 4 A). The overlap between Shot and Moe was maintained until late st.15 (Fig. 4 B). Then, we examined Shot localization in relation to MTs. Shot was strongly detected in the TC from early stages of lumen extension and until the end of TC elongation (Fig. 4 C-E). At the beginning of *de novo* lumen formation, when MTs emanated from the junction/centrosome pair, Shot co-localized with the first sprouting stable MTs (Fig. 4 C-E). The overlap between Shot and stable MTs was strongly observed also at embryonic st. 15 when a MT track preceded subcellular lumen detection (Fig. 4 C). At st. 16, both Shot and stable MTs localized to the apical side of the TCs in the area surrounding the subcellular lumen (Fig. 4 F).

**Figure 4.**
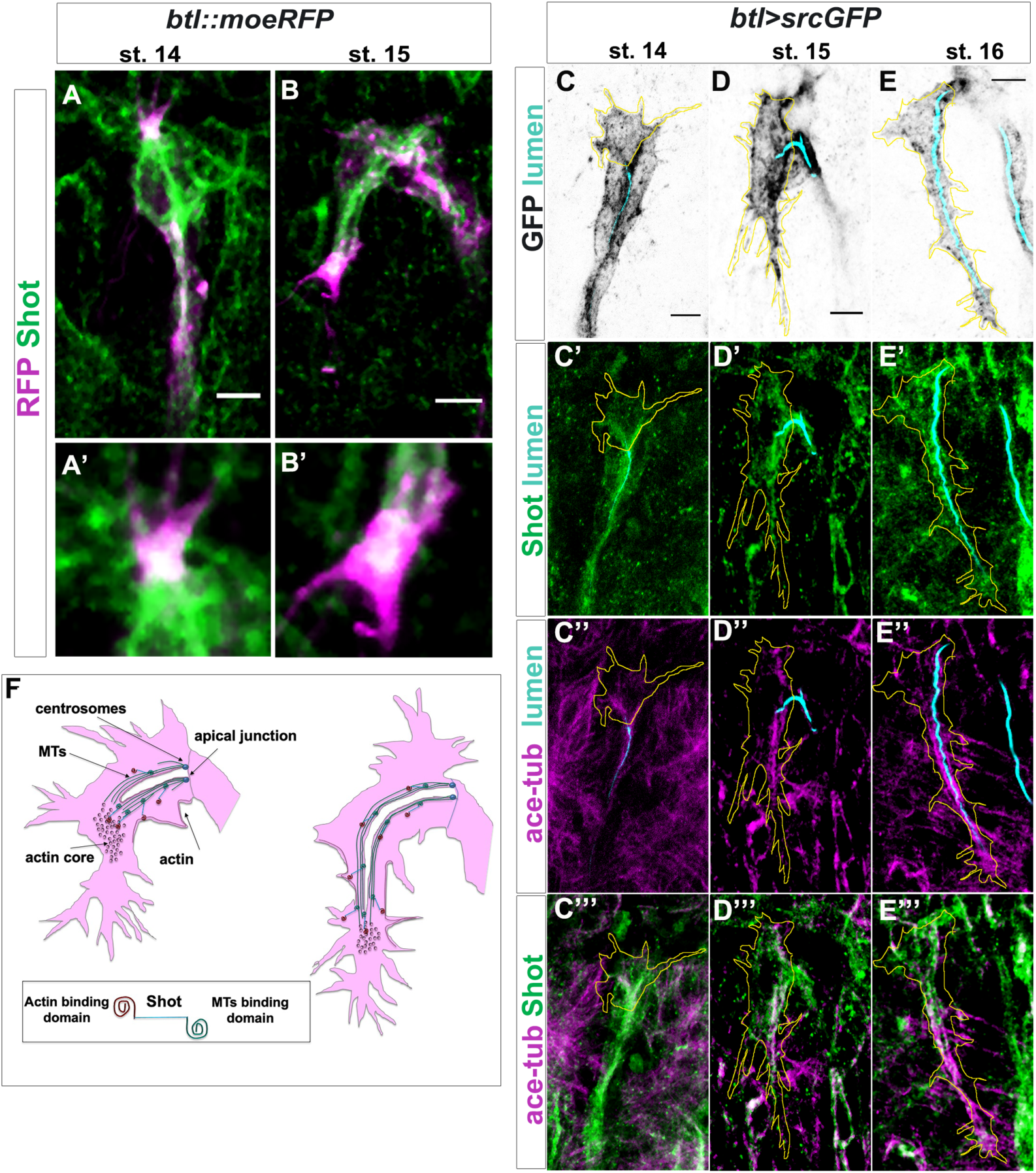
Shot colocalizes with TC cytoskeletal components. (A-B) Endogenous Shot colocalized to the Actin/Moe area during TC development. Tip of dorsal branches from st. 14 to late st. 15 of *btl::moeRFP* embryos stained with RFP (magenta) and Shot (green). In the magnification of the tip of the TCs (A’-B’) note Shot and RFP co-localization. (C-E) Endogenous Shot accumulated around stable microtubules during subcellular lumen formation. Dorsal TCs from fixed embryos *btl>srcGFP* stained with Shot and acetylated-tubulin antibodies and fluostain, from st.14 to st. 16. In all panels the TC outline is drawn in yellow. GFP staining is showed in grey and cell contour in yellow (C-E), endogenous Shot is in green (C’-E’ and C’’’-E’’’), acetylated tubulin is in magenta (C’’-E’’’) and the lumen was detected with fluostain, represented in cyan in (C’-E’ and C’’-E’’). Acetylated tubulin and Shot are both accumulated toward ahead of the subcellular lumen at earliest stages (st. 14-15) and around the subcellular lumen at later stages (st. 16). Note that co-localization between acetylated tubulin and Shot is mainly detectable inside the TCs. Anterior side is on the left, dorsal midline is up. Scale bar 5μm. (F) Schematic representation of dorsal TC development from st.15 to st.16. Basal membrane in grey, apical membrane in light blue, subcellular lumen in white, the actin network in red and MTs are in green. Between st.14 and st.15 actin dots mature in an actin core in front of the tip of the subcellular lumen in formation that is surrounded by microtubules. Shot (represented on the bottom of the figure) is detectable both inside the actin core and surrounding the lumen where stable MTs are organized.

Shot localization within the TC suggested that the spectraplakin localized with stable MTs all around the nascent lumen and with the Moe/Actin at the tip of the TC, during the time of cell elongation and subcellular lumen formation. This suggests that Shot mediates the crosstalk between these two cytoskeletal components, helping their stabilization and organisation during subcellular lumen formation and growth (Fig. 4 F).

### Absence of Shot leads to disorganized microtubules and actin

We then asked how actin and MTs were localized and organized in *shot^3^* mutant embryos. We analysed the different types of TC mutant phenotypes ranging from cases in which the TC did not elongate and the subcellular lumen was not formed, to cases in which the TC was able to elongate and form the lumen albeit not to *wt* levels (Fig. 5). In all cases, we found defects in both MTs and Moe accumulation in mutant TCs.

**Figure 5.**
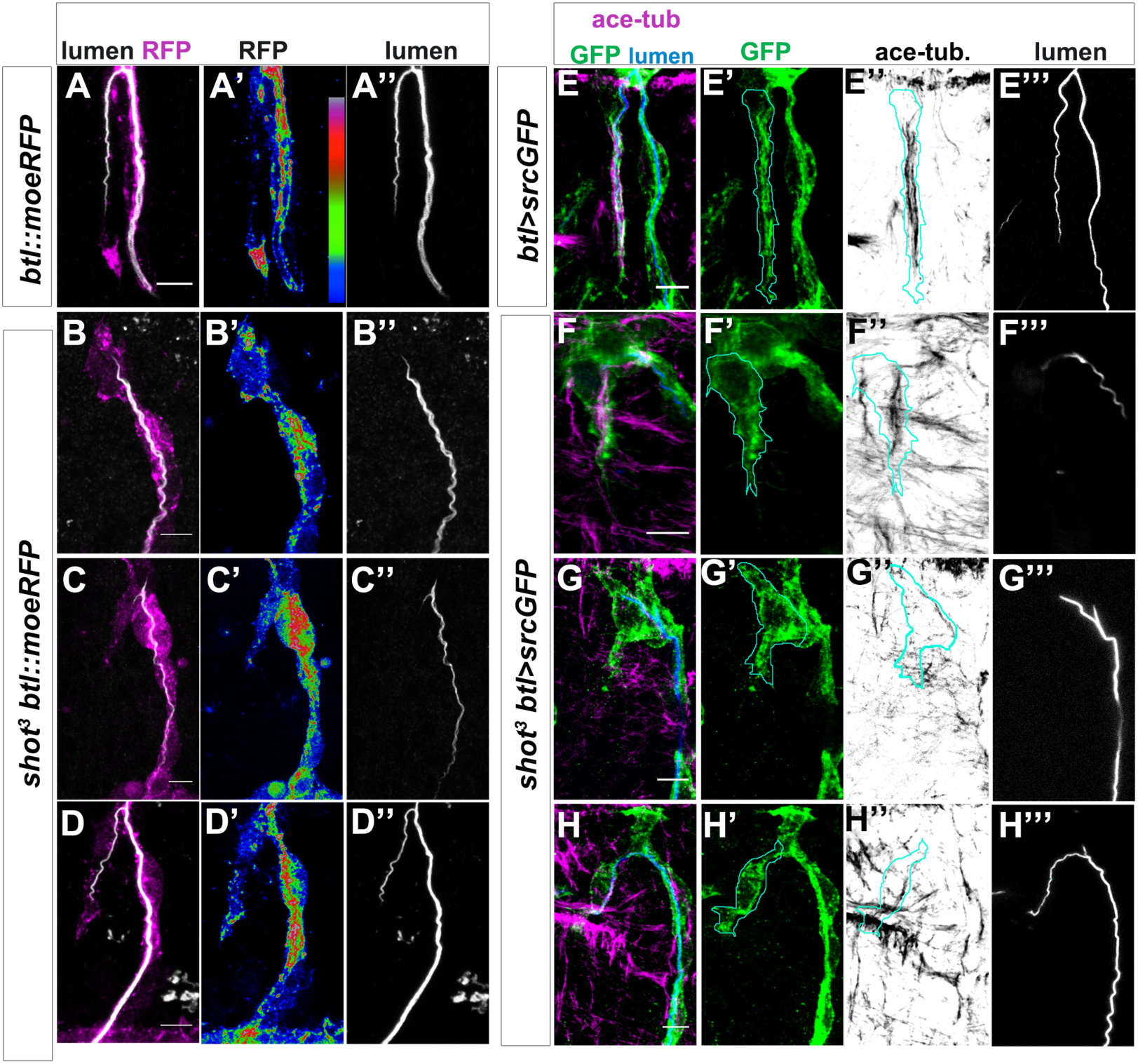
Shot LOF leads to disorganized MT-bundles and actin localization defects. (A-D) Asymmetric actin accumulation is affected in *shot^3^* mutant embryos. Dorsal TC from *shot^3^/+*; *btl::moeRFP* heterozygous controls (A) and *shot^3^*; *btl::moeRFP* mutant embryos (B-D), stained with RFP (Magenta in A-D or in a colour scale in which blue is low, green is middle and red high intensity in A’-D’) and CBP (in white). In *shot* mutant Moe/Actin was affected in its accumulation in the TC (B-D). (A) wt control; (B) when the cell was not elongated and the lumen is not formed; (C) when the cell was partially elongated but the lumen was not; (D) when the cell elongated and a lumen was detected (D). Note that Moe/Actin was affected even when the cell was elongated and a partially lumen was formed. (E-H) TC MT-bundles in *shot^3^*. Dorsal TC from a st. 16 control embryo (A) and *shot^3^* mutant (B-D) stained with GFP (green) acetylated tubulin (in magenta in E-H and in grey in E’’-H’’) and CBP (in blue in E-H and grey in E’’’-H’’’). The TC border is drawn in cyan (E-H’’). In all cases the organization and the overall amount of stable MTs detected was strongly affected; in (F) MT-bundles were observed to be disorganized along the cytoplasm devoid of a subcellular lumen and in G and H only a thin MT-bundle surrounded the subcellular lumen. Anterior side is on the left and dorsal midline is up. Scale bars 5 μm.

Considering actin localization, in control embryos at early st. 16, Moe was strongly localized at the tip of the TC, in front of the tip of the growing lumen (86% of TCs analysed, n=21). Moreover, a few spots of Moe were detectable in the cytoplasm, around the subcellular lumen (Fig.5 A and Movie 2). In *shot^3^*, we observed reduced Moe accumulation at the TC-tip and an increase of scattered spots into the cytoplasm (86% of TCs analysed, n= 23) (Fig. 5 B-D), indicating that Shot contributed to TC actin organization.

Regarding MT-bundles, we observed stable MTs organized in longitudinal bundles around the subcellular lumen in control TCs (Fig. 5 E). In *shot^3^* TCs (n=20), we detected MT-bundle defects. In particular, we observed that when the TC was not elongated, MT bundles no longer localized to the apical region and seemed to be fewer than in *wt* (Fig. 5 F). A general disorganization in MT bundles in respect to the control was also observed in TCs partially able to elongate a subcellular lumen (Fig. 5 G, H).

These analyses, taken together with the previous analysis of Shot localization in *wt* TCs, suggested a spectraplakin role in organizing/stabilizing both MTs and Moe/Actin accumulation in the TC.

### Subcellular branching depends on both actin and microtubule binding domains of Shot

In order to analyse how the different domains of Shot affected luminal development and branching, we expressed different isoforms of Shot in *shot^3^* mutant TCs. As described previously, *shot^3^* embryos displayed a variable expressivity in TC phenotypes. To simplify the quantification of the rescue experiments, we took in consideration the most severe luminal phenotype: the complete absence of a subcellular lumen. In *shot^3^*, we quantified that 22% of TCs (at the tip of GBs and DBs) did not develop a subcellular lumen at all (Fig. 1 L). Targeted expression of full-length ShotA in the trachea of *shot^3^* mutant embryos was able to rescue the subcellular lumen phenotype to the level of only 6% of the TCs analysed (n=200) not developing a subcellular lumen (Fig. 6 C).

**Figure 6.**
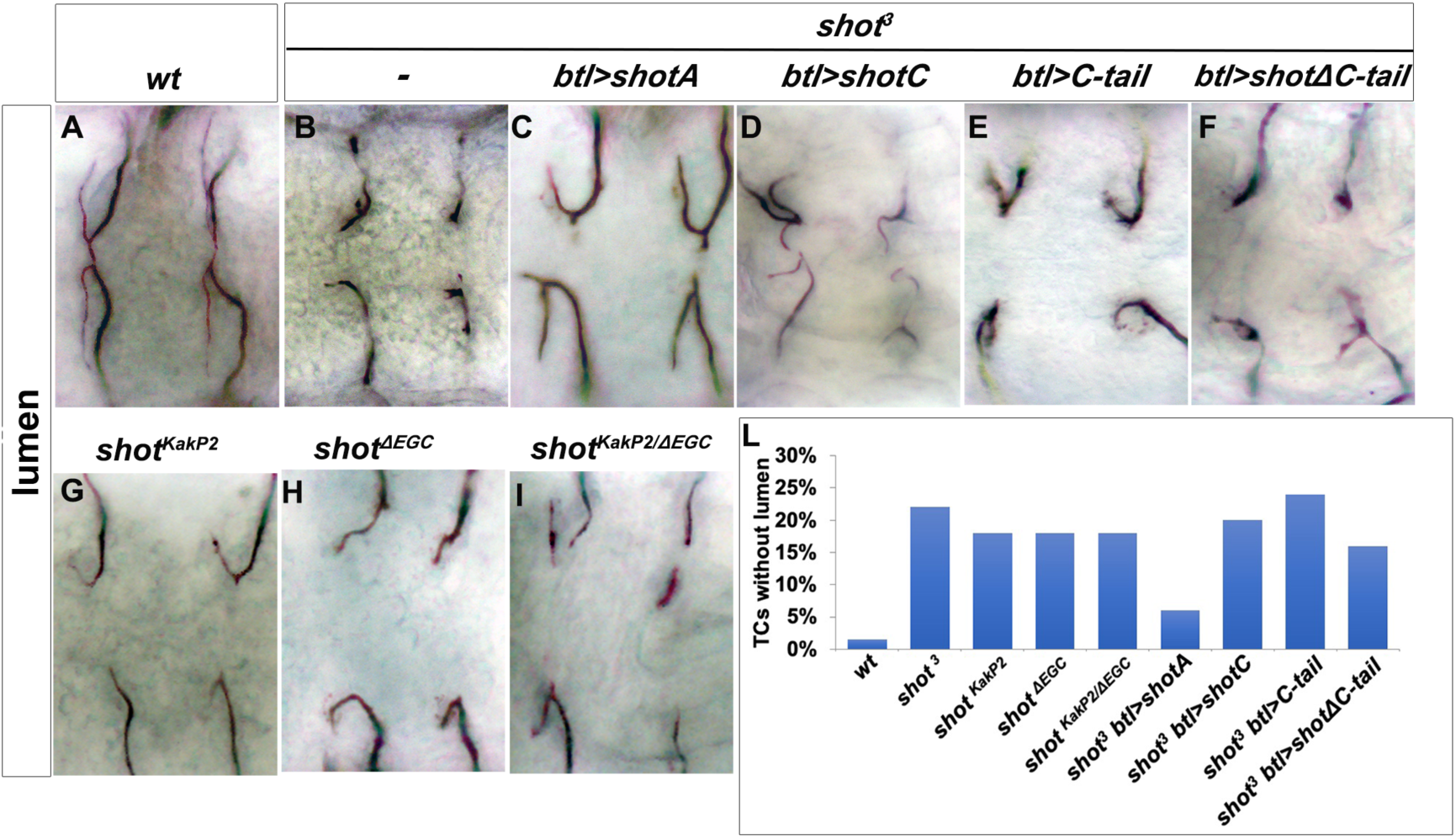
Shot Actin- and MT-binding domains are necessary for proper subcellular lumen formation. (A-I) Dorsal branches of st.16 embryos, stained with anti-Gasp to visualize the lumen. Genotype is indicated above each panel. (B-F) Null allele, *shot^3^*, rescue experiments indicate that both the actin-binding domain and the MT-binding domain are involved in subcellular lumen formation since the only construct able to rescue the null allele phenotype was full length UASShotA (B and J). (G-I) Both functional domains are needed in the same molecule since mutants affected only in the actin-binding domain (*shot^KakP2^*) or in the MT-binding domain (*shot^ΔEGD^*) and the transheterozygous *shot ^KakP2/ΔEGD^* display the same subcellular lumen phenotype as the null mutant *shot^3^.* (J) Quantification of TCs without lumen: *wt*, *shot^3^*, *shot^KakP2^*, *shot^ΔEGD^*, *shot ^KakP2/ΔEGD^*(n=400 TCs), *shot^3^*, *btl>shotA* (n=200 TCs), *shot^3^*; *btl>shotC*, *shot^3^*; btl> *shotCtail*(n=240), *shot^3^*; *btl> shotΔCtail* (n=220).

We then proceeded to molecular dissect the function of Shot in TCs. To do so, we used the three different constructs Shot: ShotC, Shot∆Ctail and ShotCtail (Fig. 2 M). When we expressed ShotC in the tracheal TCs we found that 20% of TCs analysed (n=200), had TCs with no lumen (Fig. 6 D), suggesting that the ABD domain is necessary for the correct *de novo* luminal morphogenesis.

We next expressed *shotC-tail* in order to address whether the Shot MT-binding domain alone could restore subcellular lumen formation. We observed that 24% of TCs analysed at the tip of GBs and DBs (n=250) were still not able to form a subcellular lumen (Fig. 6 E), suggesting that the tracheal expression of *shotC-tail* was not enough to rescue the null phenotype. Finally, we expressed *shot∆C-tail* to test whether Shot without the MT-binding domain could restore subcellular lumen formation. We observed that 16% of TCs analysed at the tip of GBs and DBs (n=250) were still unable to form a subcellular lumen (Fig. 6 F). Taken together, these analyses suggested that full-length isoform A, allowing Actin-MT crosslinking is necessary for correct *de novo* subcellular lumen formation.

In order to further test the hypothesis that full-length Shot is needed to correctly form a subcellular lumen, we analysed *shot^kakP2^* mutant phenotype. This allele carries an insertion of a transposable element into the intron between the second and the third transcriptional start site of *shot* abolishing all isoforms containing the first Calponin domain (CH1) and interfering with Shot actin-binding activity (33). The penetrance and expressivity of the phenotype observed in *shot^kakP2^* TCs was very similar to *shot^3^* null allele with 18% of these (n=600; 300 ganglionic and 300 dorsal TCs) not forming a subcellular lumen at all (Fig. 6 G). In addition, *shot^kakP2^* TCs displayed the same MT and actin disorganization phenotypes as *shot^3^* TCs (Fig. S4). Phenotypic data from *shot^kakP2^* together with data from transgenic rescues with the *ShotC* construct, lacking the CH1 domain, indicate that Shot full-length is required for *de novo* subcellular lumen formation. Since the actin and MT binding domains were shown to be necessary for the proper formation of a subcellular lumen, we asked whether it was necessary to have both domains in the same protein or if simply the independent presence of these domains was enough to generate a subcellular lumen. To do so, we generated transheterozygous flies expressing two different Shot isoforms, Shot^kakP2^ and Shot^∆EGC^. Shot^∆EGC^ is a truncated protein, lacking the EF-hand, the Gas2 and the C-tail domains of Shot, leading to complete loss of the MT-binding activity (38). The analysis of *shot^∆EGC^* mutant TC phenotypes revealed that 18% of TCs (n=400; 200 ganglionic and 200 dorsal TCs) did not develop a TC lumen at all (Fig. 6 H) and that *shot^∆EGC^* mutant TCs displayed MT and actin disorganization phenotypes (Fig. S4). Interestingly, in *shot^∆EGC^* mutant TCs, actin was found to be disorganized throughout the cytoplasm (and not at the tip as in control TCs) but in higher levels than in *shot^kakP2^* TCs (Fig. S4). This suggests that the actin-binding domain present in *shot^∆EGC^* is able to organize the actin in TCs albeit not to wt levels.

In *shot^∆EGC^*/*shot^kakP2^* transheterozygous embryos, Shot molecules contained exclusively either the CH1 or the C-tail, but neither molecule had actin- and MT-binding activity simultaneously. These embryos displayed the same TC phenotype as either homozygous mutant (18% TCs with no lumen, n=400) (Fig. 6 I), indicating that both the actin- and the MT-binding domains need to be present in the same Shot molecule for proper TC subcellular lumen formation.

Taken together these results indicate that Shot is able to mediate the crosstalk between MTs and actin during subcellular lumen formation, via its MT and actin-binding domains and that these have to be present in the same molecule for proper subcellular lumen formation.

### Increased levels of Shot are induced in TCs by DSRF

The TC-specific transcription factor *bs*/DSRF is important for TC specification and growth, and has been suggested to regulate the transcription of genes that modify the cytoskeleton (9, 39). Considering the luminal phenotypes associated with *bs* LOF in TCs and the role of MTs in subcellular luminal formation, we asked whether *shot* expression in TCs could be regulated by DSRF.

In order to test this, we searched *in silico* for DSRF binding sites in the promoter regions of all *shot* isoforms using the Matscan software (40) and the reported position weight matrix (PWM) corresponding to SRF (41) (Supplementary Table 2). We found 7 regions with at least one putative binding site (binding score larger than 70% of maximum value) within 2000 bases of the *shot* annotated TSS (Fig. 7 D and Supplementary Table 1). These regions mapped to the locations of known Shot promoters (Fig. 7 D) (36). We then asked if lower Shot levels could be detected in *bs* mutant TCs. Indeed, when analysing *bs* in comparison to *wt* TCs, we could detect lower levels of endogenous Shot protein (Fig. 7 A-C). To confirm this, we analysed the TC phenotype of *bs* embryos upon tracheal expression of *shot* in these cells. We observed that increasing *shot* expression in TCs resulted in rescue of *de nov*o lumen formation in *bs* TCs (Fig. 7 E-K). Taken together these results indicate that at least part of the luminal phenotypes associated with *bs* LOF in TCs are due to lower levels of the Actin/MT binding activity of Shot.

**Figure 7.**
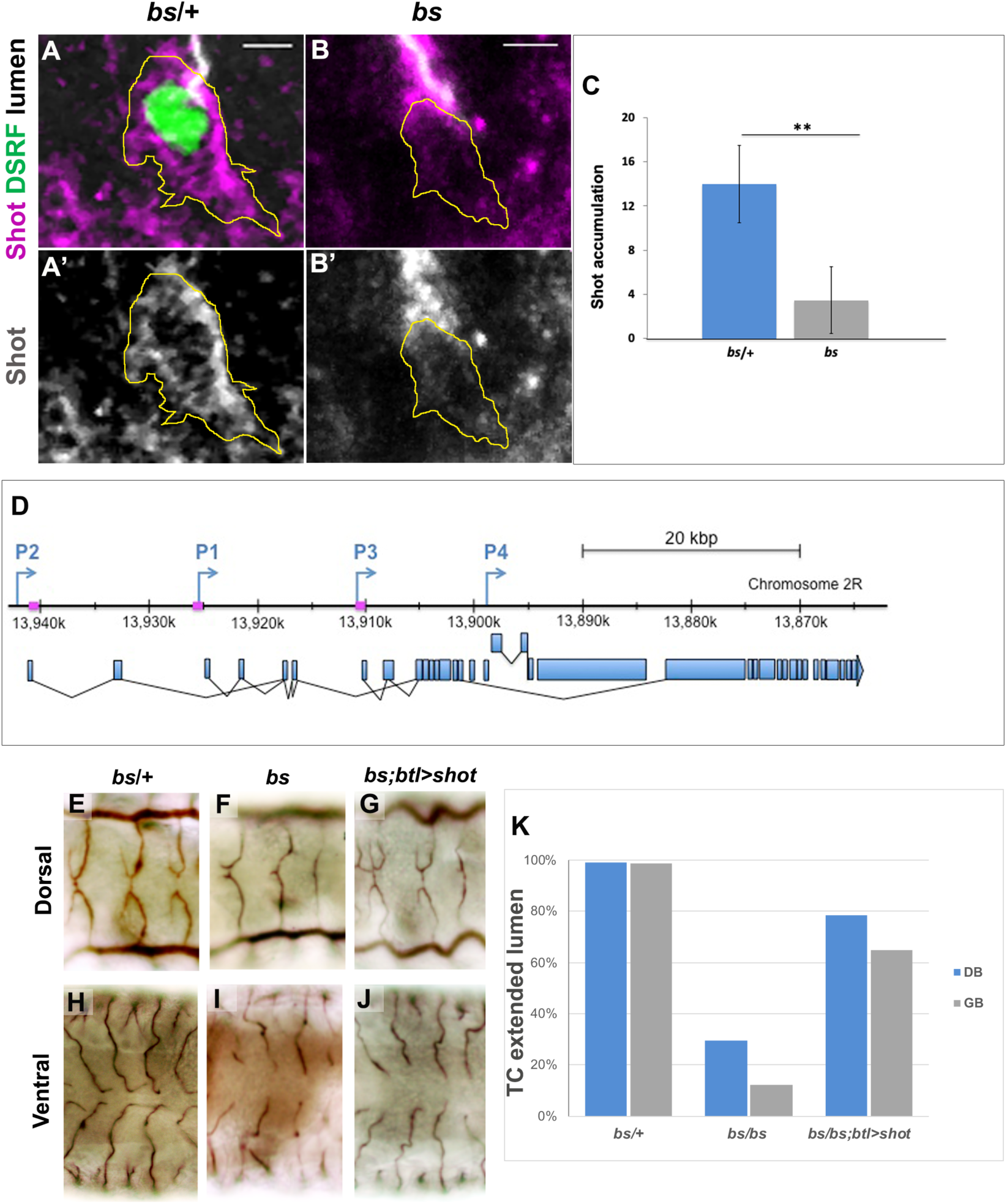
Shot expression is regulated by DSRF in TCs. GB TC at st. 15 from *bs* heterozygous controls (A) and homozygous mutant embryos(B), stained with Shot (magenta in A and B, grey in A’ and B’), DSRF (green) antibodies and CBP (grey). In yellow, the outline of the TCs. Shot is less accumulated in TCs from homozygous (B, B’ and C) n=9 TCs (raw integrated density was measured +/− SEM). (D) P1, P2 and P3 transcription start sites of the *shot* locus together with the specific sequences recognized by the DSRF transcription factor (squares in magenta) (adapted from (36)). Dorsal and ventral TCs from *wt* (E and H) *bs* (F and I) mutant embryos. The tracheal overexpression of *Shot* is sufficient to restore the growth of TC subcellular lumina in *bs* mutant background (G, J). (K) Quantification of TCs with an extended lumen: *bs*/+ n=350; *bs*/*bs* n=280 and *bs*/*bs*;btl>ShotC n=210.

### Shot and Tau functionally overlap during subcellular lumen formation and branching

Previous *Drosophila* work suggested that Shot could display potential functional overlap with Tau in microtubule stabilisation (35, 42). To assess this functional overlap during TC subcellular branching, we started by overexpressing Tau in TCs using GAL4 induced expression. Upon overexpression of Tau in otherwise *wt* TCs, we detected ESLs in 93% of TCs, which is comparable to the ShotOE phenotype (Fig. 8 A-C). Like in ShotOE, this effect was dosage dependent, with more TCs with ESLs when more Tau copies were expressed (Fig. 8 C). We then tried to rescue the *shot* LOF phenotype by targeted expression of Tau in TCs. Again, this effect was dosage dependent. We achieved a 64% rescue of the *shot* mutant phenotype with two copies of Tau expressed, indicating that Tau can execute a similar function to Shot in *de novo* subcellular lumen formation (Fig. 8 D-L).

**Figure 8.**
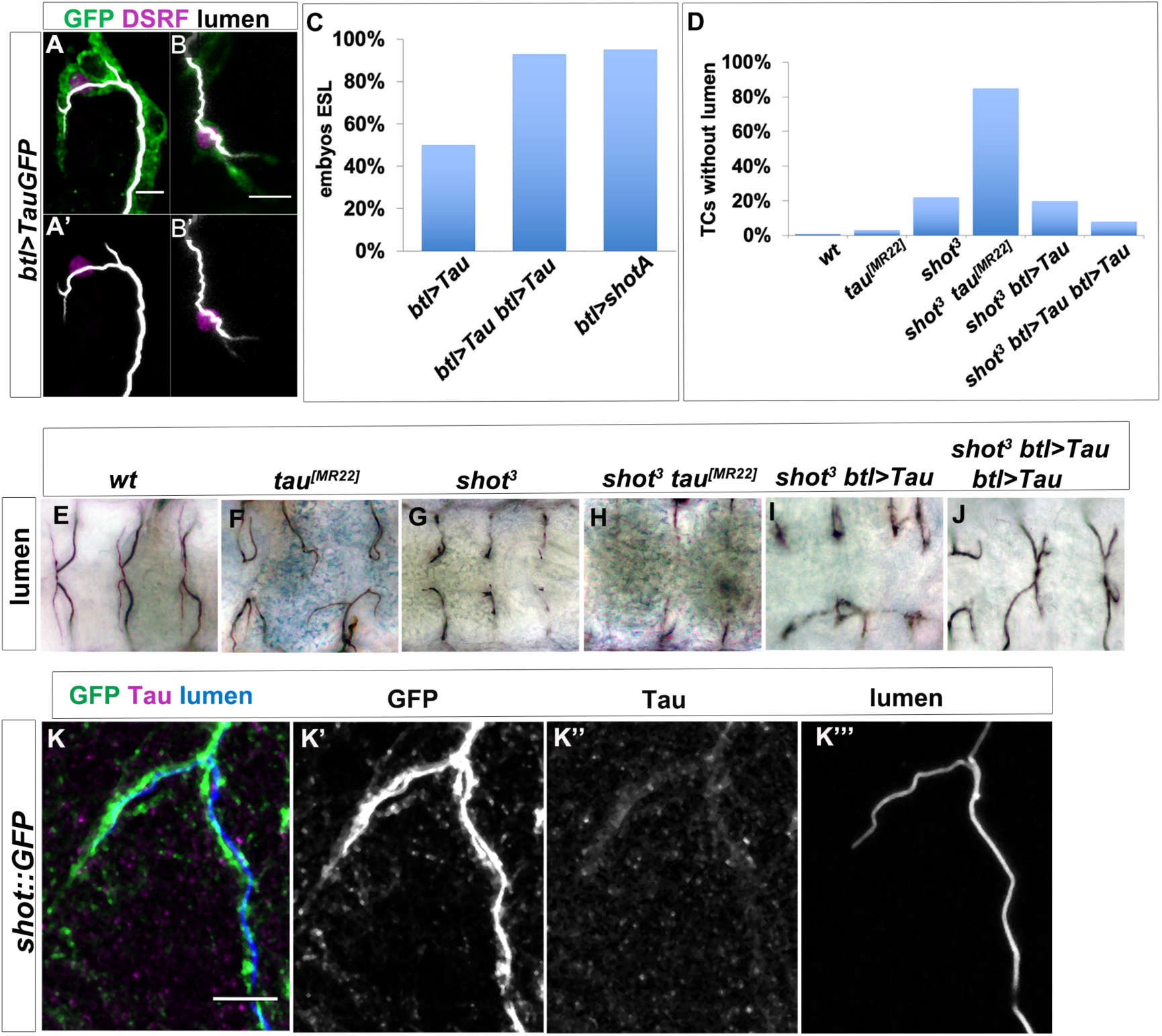
Shot and Tau functionally overlap during subcellular lumen formation. (A-B) DB (A) and GB (B) embryonic TCs expressing *TauGFP* in the tracheal system, stained with GFP (green), CBP (white) and DSRF (magenta), showing the ESL phenotype. In A’ in B’ lumen and TC nuclei are shown, anterior side on the left, dorsal side is up; scale bar 5 µm. (C) Quantification of embryos overexpressing one (n=23) or two (n=16) copies of *btl > TauGFP* displaying at least one bifurcated terminal cell in comparison with the ShotOE quantification (n= 20). (D) Quantification of TCs without subcellular lumen and (E-J) dorsal view of TCs from st. 16 embryos (genotype indicated) stained with anti-Gasp. *tau* deletion mutant does not display a subcellular lumen phenotype (D and F, nTCs =79) but enhances the effect of *shot* mutation in the double mutant *shot^3^; tau^[MR22]^* (D and H, TCs=180). One copy of Tau is not sufficient to rescue *shot^3^* (D and I, n=400) but two copies rescues the *shot* LOF TC phenotype (D and J n=260). (K) Tau is detected in embryonic TCs. Embryonic *shot::GFP* dorsal TC stained with GFP (green in K, grey in L), anti-Tau antibody (magenta in K, grey in M) and CBP (blue in K grey in N); scale bar 5 µm.

We then asked whether *tau* null mutants displayed any TC luminal phenotypes. For this we analysed a *tau* deletion mutant *tau^MR22^* previously shown to have nervous system defects (42). *tau^MR22^* null mutant TCs showed defects in subcellular lumen directionality, but not in subcellular lumen formation (Fig. 8 D, F). We then proceeded to analyse TCs in double mutants for *shot^3^* and *tau^MR22^* (*shot-tau*). These double mutants showed higher numbers of TCs without lumen (85%) than TCs from *shot^3^* (22%) or *tau ^MR22^* (3%) alone, or a mere sum of these phenotypes, indicating a synergistic genetic effect between *shot* and *tau* (Fig. 8 D-H). Despite the strong phenotypes, *shot-tau* mutants have the correct number of cells per branch and express DSRF in TCs (Fig. S5). Furthermore, using a mouse anti-Tau antibody, we could detect Tau colocalizing with the growing lumen in TCs (Fig. 8 K). These results indicate that, as seen in neurons (42), in tracheal TCs Shot and Tau functionally overlap in subcellular lumen formation and branching.

### Shot is required for subcellular luminal branching at larval stages

During larval stages, TCs ramify extensively to form many branches from the same cell body, long cytoplasmic extensions that form one cytoplasmatic membrane-bound lumen each (3, 16). We questioned if Shot was also necessary for the subcellular branching and lumen extension in these larval cells. To answer this, we expressed different isoforms of Shot, Shot-RNAi and Tau in TCs from embryonic stages with a TC specific driver (DSRF-GAL4) and analysed the phenotypes on branching and ESL formation at the end of the larval stages (Fig. 9). Downregulation of Shot induced TCs with lower levels of branching and fewer lumina (Fig. 9 B, G, I). Whereas in the *wt* each TC branch is filled by a subcellular lumen, in Shot-RNAi TCs these were reduced to 37% of the TCs and even so absent in most branches (Fig. 9 B and G). Also, on average, each *wt* TC develops 17 branch points, but Shot-RNAi TCs only developed an average of 6,5 branch points each (n=8) (Fig. 9 B and I). We then overexpressed Shot full-length (ShotA-GFP aka ShotOE condition) and could not detect extra branching points in TCs, suggesting that more than just an increased Actin-MT crosstalk is needed for the induction of TCs with supernumerary cytoplasmatic extensions. Nonetheless, overexpression of ShotA, ShotCtail and Tau induced ESL in TCs, with 2 or more lumina in all cells analysed (n=10) (Fig. 9 C-E and H). Like in embryos, targeted expression of Shot-∆C-tail did not induce ESL in larval TCs (Fig. 9 F and H). Taken together, these results indicate that Shot is necessary for larval lumen formation and branching and that Actin-MT crosstalk by Shot or Tau is sufficient for ESL formation within each TC cytoplasmatic extension.

**Figure 9.**
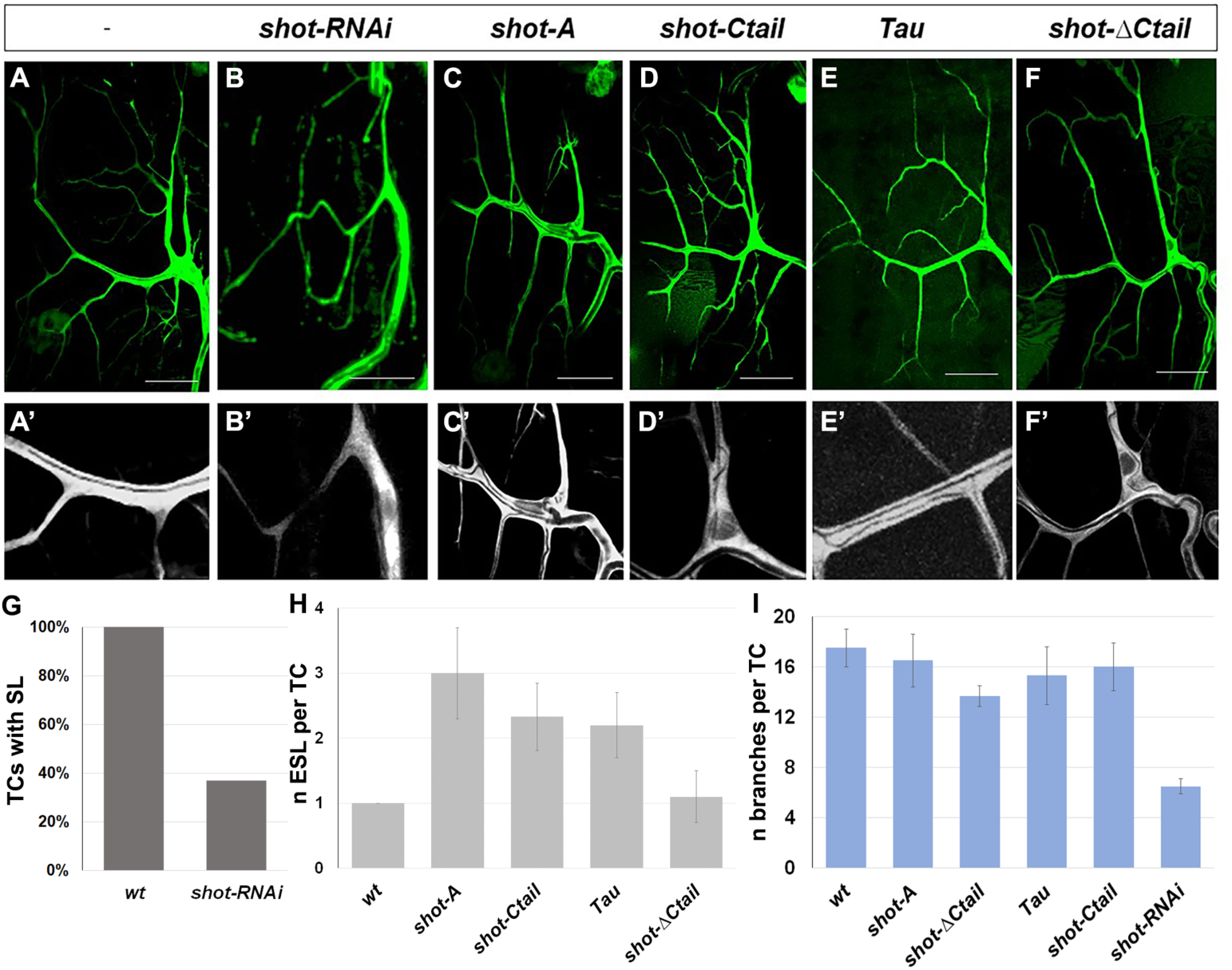
Shot and Tau modulate luminal branching in larval TCs. Wandering larval (L3) TCs expressing only GFP (A) and different Shot and Tau constructs (B-F) under the control of a tracheal DSRFGAL4 driver (all except E where the driver used was btlGAL4). (A, A’) UASEB1GFP (n=10) (B, B’) UASshotRNAi and UASEB1GFP (n=10); (C, C’) UASShotA-GFP (n=10); (D, D’) UASshotCtail-GFP (n=8); (E, E’) UAStauGFP (n=8); (F,F’) UASshot∆Ctail-GFP (n=8). (G) Quantification of the percentage of TCs with detectable subcellular lumen; (H) quantification of the number of ESL per TC branch found; (I) quantification of the number of branches per TC. Scale bars 50 μm.

## DISCUSSION

In this study, we analysed the importance of MT-actin crosstalk through Shot and Tau in subcellular lumen formation in *Drosophila* embryonic and larval tracheal cells. Our work reveals novel insights into the formation of lumina by single-cells. First, that a spectraplakin in involved in the crosstalk between actin and MTs in tracheal TCs and that this crosstalk is necessary for *de novo* lumen formation. Absence of Shot leads to defects in microtubule and actin organization and a profound alteration of the cytoskeleton in TCs (Fig. 10 A, C). Second, that once a primary lumen is formed *de novo* in TCs, neither actin-MT crosstalk, nor supernumerary centrosomes, are necessary for the formation of new supernumerary lumina (ESLs). New lumina can arise from branching points along the length of the pre-existing lumen, only by MT stabilisation (Fig. 10 B, D). In these cases, we can form ESLs acentrosomally, probably from the MTOC activity provided by the gamma-tubulin present along the crescent lumen (1). Third, spectraplakin activity is necessary to organize MTs and actin in TCs. Fourth, increased levels of Shot are induced in TCs by DSRF, and Shot can rescue the subcellular lumen formation phenotypes in *bs* mutants. This agrees with previous observations in other systems where *bs* and *shot* mutants display similar phenotypes (43). And fifth, high-levels of Tau can replace Shot in subcellular lumen formation and branching.

**Figure 10.**
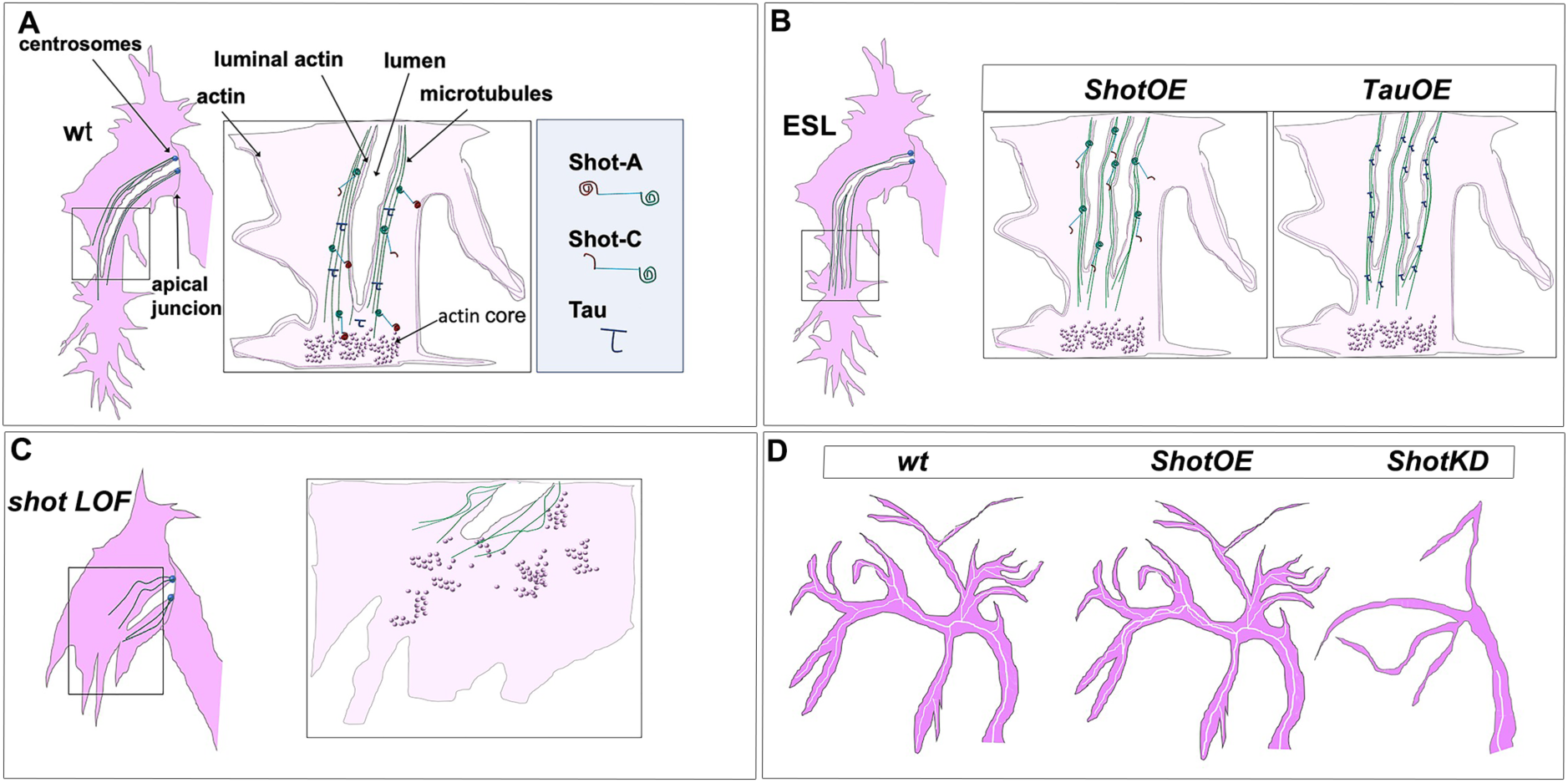
Shot and Tau dynamically modulate the cytoskeleton during subcellular lumen formation. Schematic representation of st.16 embryonic (A, B, C) and third instar larval (D) TCs; cytoplasm is in pink and luminal space in white. (A) Cytoskeletal components in a *wt* embryo with the actin-network (dark pink) and MTs (green). Shot and Tau are able to organize the cytoskeleton by crosslinking MTs and actin; Shot (represented with the actin domain in red and the MT binding domain in green) mediates the crosstalk between actin and MTs as the longer isoform (ShotA), but shorter isoforms lacking the ABD (ShotC) do not bind actin. Tau Is represented in blue. (B) ESLs are formed by from the pre-existing lumen, which acts as an MTOC, by overexpressing *shot* (ShotOE) or *Tau* (TauOE) through MT-stabilization. (C) In the absence of Shot proper cytoskeletal organization is not established and cell elongation and lumen formation do not occur. This phenotype can only be rescued by expressing full-length Shot in TCs. (D) Schematic representation of larval TCs in *wt*, in *ShotOE* (or *TauOE*) where ESLs are formed without concomitant single-cell branching and in *Shot*KD, where both cellular and luminal branching are reduced.

### Shot promotes subcellular branching by organizing and mediating the crosstalk between microtubules and actin

Previously, it was shown that Shot was involved in tracheal fusion cell anastomosis during embryonic development (25). It was observed that Shot accumulates at E-cadherin-dependent contacts between fusion cells and *shot* LOF disrupts this contact leading to cell-fusion phenotypes. In these cells, interactions of Shot with F-actin and microtubules are functionally redundant and both targeted expression of ShotC or ShotA is sufficient to rescue the cell-fusion phenotype (25). Our results are more akin to what has been reported in neuronal growth cones, and both actin and MT binding domains of Shot are required for TC extension and subcellular lumen formation (Fig. 10 A). In neurons, like in tracheal cells, ShotC is unable to rescue the phenotype caused by *shot* LOF, which is only rescued by expression of the full-length ShotA isoform (24). Shot has also been shown to be required for sealing epithelial sheets during dorsal closure (38). In these epithelial cells, Shot acts as a MT-actin crosslinker to regulate proper formation of the MT network. As in the case of tracheal TCs presented here, the actin- and microtubule-biding activities of Shot are simultaneously required in the same molecule, indicating that like in TCs Shot is engaged as a physical crosslinker also during dorsal closure (38).

MTs and the actin cytoskeleton perform many functions in tracheal TCs that are regulated by different actin- and MT-binding proteins. While mediators of actin function, such as Ena (1), and of MT function, like D-Lissencephaly-1 (DLis-1), have been identified previously, we show here that Shot is able to crosstalk MTs and actin during subcellular lumen formation. In Shot LOF conditions, MTs and actin are disorganized. Consequently, this Shot crosslinking function is essential for *de novo* lumen formation and extension. It has been previously described that in TCs of mutants affected in MT organization, the actin-network is not perturbed (1), so the “actin phenotype” observed in *shot* LOF cannot be a consequence of defects in the MT network. This observation indicates a possible spectraplakin function in organizing TC actin in agreement with previous observations that Shot and ACF7 can promote filopodia formation (37, 44).

### Shot expression is regulated by DSRF in TCs

Our results show that molecular levels of Shot are important for cytoskeletal rearrangements, indicating that there is a dosage dependent effect in lumen formation and extension as well as in luminal branching events. Shot is present in many cells during development but Shot level regulation is likely to be more important in cells such as neurons and tracheal terminal cells, due to their morphology (34). *bs*/DSRF is a TC-specific transcription factor, whose expression is triggered by Bnl signalling (9, 45), and is required for TC cytoskeletal organisation (1). DSRF has also been shown to be necessary not just for the establishment of TC fate, but to ensure the progression of TC elongation (32). Cytoskeletal organisation and remodelling as well as TC elongation are tightly coupled during subcellular lumen formation and in *bs* mutants actin accumulation was impaired at the TC tip (1). We observe a similar actin phenotype in Shot mutants (Fig. 5 A-D) suggesting that the actin defects observed in DSRF mutants may be due to a lower expression of Shot in these cells.

### Shot and Tau functionally overlap in subcellular lumen formation and branching

It has been suggested that spectraplakins functionally overlap with structural microtubule-associated-proteins (MAPs). Shot displays a strong functional overlap with Tau in MT stabilization leading to the adequate delivery of synaptic proteins in *Drosophila* axons (42). In addition, it has been proposed that a loss of MAP function in mammals results in a relatively mild phenotype due to a functional compensation accomplished by spectraplakins (46, 47). Furthermore, the effect of the complete lack of Shot function during dorsal closure is very subtle (38), hinting that in another *Drosophila* organ, Shot function might have overlaps with other MAPs.

Our overexpression and genetic data suggest that also in the context of subcellular lumen formation these two proteins functionally overlap. When we tested the tracheal overexpression of *tau* in *wt* background, we observed extra subcellular lumina with morphology very similar to the one caused by ShotOE. Moreover, Tau overexpression in tracheal cells was able to rescue the *shot* LOF phenotype similarly to ShotA expression. We propose that Tau’s rescuing capability does not depend only on its classical MT-stabilization activity, since expression of ShotC and ShotC-tail in tracheal cells was not able to restore subcellular lumen formation. Tau MT-binding is probably just one of its functions in TCs. In fact, Tau has been show to co-organize dynamic MTs and the actin-network in cell-free systems and growth cones (48–50). Our rescue and double mutant analyses suggest that in TCs, Shot and Tau functionally overlap in organizing the coordination between MT-bundling and actin cytoskeleton crosstalk (Fig. 10 A, B).

### Larval lumen formation and branching

TC subcellular lumen formation starts at embryonic stages but most of its elongation and branching occurs during the extensive body growth of the third instar larva (L3). Some mutants have been reported to generate larger TCs with higher numbers of branches. Such mutants included the Hippo pathway member *warts/lats1* (aka *miracle-gro*), and the TOR pathway inhibitor, *Tsc1* (aka *jolly green giant*) (16). In addition, activation of the FGF Receptor (Btl) pathway in TCs gives rise to ectopic branches (17, 51). Interestingly, in all these cases, mutant TCs develop a higher number of branches but no reported ESL per branch. In larvae, as in embryonic TCs, actin is present at the basal plasma membrane and at the luminal/apical membrane. The connection between the basal actin network and the outer plasma membrane is made through Talin, which links the network to the extracellular matrix (ECM) via the integrin complex (52). Regulation of the luminal actin is done by Bitesize (Btsz), a Moe interacting protein (13). These interactions with actin are required for proper TC morphology, and mutations in either the *Drosophila* Talin gene *rhea* or *btsz* induce multiple convoluted lumina per TC branch (13, 52). *rhea* and *btsz* ESLs seem to be misguided within the TC and present a series of U-turns and loops we did not observe in *shot* mutants. Also, mutations in *rhea* and *btsz* do not induce embryonic TC luminal phenotypes, suggesting that despite their interactions with actin the mechanism of action during subcellular lumen formation and stabilization is different.

They do not seem to interact with MTs and they might have a more structural/less dynamic role in larval subcellular lumen formation. Our results suggest that Shot is able to induce larval ESLs by the same mechanism as in embryos. By modulating a dynamic crosstalk between MTs and actin that induces acentrosomal luminal branching. However, albeit necessary for larval luminal branching excess Shot alone is not sufficient to induce extra branching in TCs. Perhaps ShotOE TCs are able branch their subcellular lumen but lack a specific spatial cue to induce single-cell branching. This cue could be such as the one provided by a hypoxic tissue secreting the FGFR ligand, Bnl, which would allow for the cytoplasmic extensions needed to increase single-cell TC branching.

### Shot and lumen formation in other organisms

The spectraplakin protein family of cytoskeletal regulators is present throughout the animal kingdom. In the most commonly studied model organisms we find VAB-10 in the worm *Caenorhabditis elegans*, and, in vertebrates, dystonin (also known as Bullous Pemphigoid Antigen 1/BPAG1) and Microtubule-Actin Crosslinking Factor 1 (MACF1; also known as Actin Crosslinking Family 7/ACF7, Macrophin, Magellan) (34). They are usually strongly expressed in the nervous system and most of their functions have been unraveled by studying nervous system development and axonal cell biology (53). Spectraplakin roles have also been reported in cell-cell adhesion and cell migration (31). Recently, attention has gone into the role of spectraplakins not only during normal cellular processes but also in human disease, from neurodegeneration to infection and cancer (53). However, not much is known about a role for spectraplakins neither during lumen formation nor during subcellular branching events. Here, we provide evidence for the involvement of the *Drosophila* spectraplakin Shot in subcellular lumen formation in branching. Through its actin- and MT-binding domains, Shot is necessary for subcellular lumen formation and branching (Fig.10). This function can be functionally replaced by Tau, another microtubule associated protein which has been shown to be able to crosslink MTs and actin (50). A similar crosslink between MTs and actin may in place during vertebrate lumen formation and in other subcellular branching events.

## MATERIALS AND METHODS

| Reagent type | Designation | Source or reference | Additional information |
| --- | --- | --- | --- |
| Genetic reagent<br>(*D. melanogaster*) | *shot*<sup>3</sup> | BDSC | Lee et al., 2000 |
| Genetic reagent<br>(*D. melanogaster*) | *shot*<sup>*kakP2*</sup> | BDSC | Gregory and Brown, 1998 |
| Genetic reagent<br>(*D. melanogaster*) | *shot*<sup>ΔEGC</sup> | F. Jankovics | Takács et al., 2017 |
| Genetic reagent<br>(*D. melanogaster*) | *Rca1*<sup>G012</sup> | S.J. Araújo | Ricolo et al., 2016 |
| Genetic reagent<br>(*D. melanogaster*) | *tau*<sup>[MR22]</sup> | BDSC | Doerflinger et al., 2003 |
| Genetic reagent<br>(*D. melanogaster*) | *btl::moeRFP* | M. Affolter | Ribeiro et al., 2004, |
| Genetic reagent<br>(*D. melanogaster*) | *btl-Gal4* | BDSC | Shiga Y., 1996 |
| Genetic reagent<br>(*D. melanogaster*) | *DSRF4X-Gal4* | A.Ghabrial | unpublished |
| Genetic reagent<br>(*D. melanogaster*) | *UAS-shot L(A)*<br>*GFP* | BDSC | Lee and Kolodziej, 2002 |
| Genetic reagent<br>(*D. melanogaster*) | *UAS-shot L(C)-*<br>*GFP* | BDSC | Lee and Kolodziej, 2002 |
| Genetic reagent<br>(*D. melanogaster*) | *UAS-shot-LA-*<br>Δ*Ctail-GFP* | N. Sanchez-Soriano | Alves-Silva et al., 2012 |
| Genetic reagent<br>(*D. melanogaster*) | *UAS-shot-LA-*<br>*Ctail-GFP* | N. Sanchez-Soriano | Alves-Silva et al., 2012 |
| Genetic reagent<br>(*D. melanogaster*) | *UAS-TauGFP* | M. Llimargas | Llimargas et al., 2004 |
| Genetic reagent<br>(*D. melanogaster*) | *UAS-srcGFP* | BDSC | Kaltschmidt et al., 2000 |
| Genetic reagent<br>(*D. melanogaster*) | *UAS-shot RNAi* | BDSC | - |
| Genetic reagent<br>(*D. melanogaster*) | *shot::GFP* | J.Pastor-Pareja | Sun T. et al., 2019 |
| Antibody | mouse<br>anti GASP | DSHB | 1:5 |
| Antibody | rat<br>anti-DEcad | DSHB | 1:100 |
| Antibody | guinea Pig<br>anti-CP309 | V. Brodou | 1:1000 |
| Antibody | rabbit and rat<br>anti-DSRF | J. Casanova | 1:500 |
| Antibody | goat, rabbit Anti<br>GFP | Roche /<br>Jackson | 1:500 |
| Antibody | chicken, rabbit,<br>mouse<br>anti-βgal | Cappel/<br>Promega/<br>Abcam | 1:500 |
| Antibody | mouse anti-<br>acetylated tubulin | Sigma | 1:100 |
| Antibody | guinea pig<br>anti-Shot | K. Röper | 1:1000 |
| Antibody | mouse anti-Tau-1 | Sigma-Aldrich | 1:200 |
| Antibody | Cy2-conjugated<br>secondary<br>antibody | Jackson<br>Labs | 1:500 |
| Antibody | Cy3-conjugated<br>secondary<br>antibody | Jackson<br>Labs | 1:500 |
| Antibody | Cy5-conjugated<br>secondary<br>antibodies | Jackson<br>Labs | 1:500 |
| Antibody | Alexa 488<br>conjugated<br>secondary<br>antibodies | Thermo<br>Fischer<br>Scientific | 1:500 |
| Antibody | Alexa 647<br>conjugated<br>secondary<br>antibodies | Thermo<br>Fischer<br>Scientific | 1:500 |
| Antibody | Alexa 555<br>conjugated<br>secondary<br>antibodies | Thermo<br>Fischer<br>Scientific | 1:500 |
| Antibody | biotinylated anti mouse IgM | Thermo Fischer Scientific | 1:500 |
| Probe | CBP | J. Casanova | 1:500 |
| Probe | Fluostain | Sigma-Aldrich | 1:300 |
| AB solution | Vectastain ABC kit | Vector Laboratories | 1:200 |

### *D. melanogaster* strains and genetics

*shot^3^* (Lee et al., 2000), *shot^kakP2^* (Gregory and Brown, 1998), *shot^ΔEGC^* (Takács et al., 2017), *Rca1^G012^* (Ricolo et al., 2016), tau^[MR22]^ (Doerflinger et al., 2003) *btl::moeRFP* (Ribeiro et al., 2004), *btl-Gal4* (Shiga Y 1996*), DSRF4x-Gal4* (gift from A. Ghabrial) *UAS-shot L(A) and GFP and UAS-shot L(C)-GFP* (Lee and Kolodziej 2002), *UAS-shot-L(A)-ΔCtail-GFP and UAS-shot-L(A)-Ctail-GFP* (Alves-Silva et al. 2012), *UAS-TauGFP* (Llimargas et al., 2004), *UAS-srcGFP* (Kaltschmidt et al., 2000), *UAS-shot RNAì* (Bloomington stock centre), *shot::GFP* (Sun., T., 2019).

Chromosomes were balanced over LacZ or GFP-labelled balancer chromosomes. Overexpression and rescue experiments were carried out either with *btl-GAL4* or *DSRF4X-GAL4 at 25°C*.

### Immunohistochemistry, image acquisition, and processing

All stage embryos, collected on agar plates overnight (O/N), were dechorionated with bleach and fixed for 20 min (or 10 min for MT staining) in 4% formaldehyde, PBS (0.1 M NaCl 10 mM phosphate buffer, pH 7.4) / Heptane 1:1. Washes were done with PBT (PBS, 0.1% Tween). Primary antibody incubation was performed in fresh PBT-BSA O/N at 4°C. Secondary antibody incubation was done in PBT-BSA at room temperature (RT) in the dark for 2h.

For DAB histochemistry (used to recognize 2A12/anti-Gasp antibody) after incubation with secondary antibody (mouse IgM biotinylated antibody) embryos were treated with AB solution for 30 min at R/T (Avidin-Biotinylated Horseradish Peroxidase H from Vectastain-ABC KIT of Vector Laboratories 1:200 in PBT).

Embryos were incubated with the DAB solution (DAB 0,12% Nickel-Sulphate-Cobalt Chloride, 0,3 %H202) until black colour was achieved, usually 2/3 min.

The primary antibodies used were: mouse anti-Gasp (2A12) 1:5, rat anti-DE-cad (DCAD2) 1:100, from Developmental Studies Hybridoma Bank (DSHB), guinea pig anti-CP3019 (from V. Brodu) 1:1000, rabbit and rat anti-DSRF 1:500 (both produced by N. Martín in J. Casanova Lab), goat and rabbit anti-GFP 1:500 (From Roche and Jackson), chicken, rabbit and mouse anti-βgal 1:500 (Cappel, Promega, Abcam), mouse anti acetylated tubulin 1:100 (Sigma), guinea pig anti-Shot 1:1000 (K. Röper), mouse anti-Tau-1 (Sigma Aldrich).

Cy3, Cy2 or Cy5 conjugated secondary antibody (Jackson Immuno Research) or Alexa 488, Alexa 647 and Alexa 555 conjugated secondary antibody (Thermo Fischer Scientific) from donkey and/or goat were used 1:500 in PBT 0,5% BSA. Two probes, to label luminal chitin were used: Fluostain 1:200 (Sigma), and chitin binding protein CBP 1:500 (produced by N. Martín in J. Casanova Lab). Bright field photographs were taken using a Nikon Eclipse 80i microscope with a 20X or 40X objective. Photoshop 21.1.1 was used for measurements, adjustments and to assemble figures. Florescence confocal images of fixed embryos where obtained with Leica TCS-SPE system using 20X and 63X (1.40-0.60 oil) objectives (Leica). Fiji (Imagej 1.47) (54) was used for measurements and adjustments. The images shown are, otherwise stated in the text, max-intensity projection of Z-stack section.

### Quantification and Statistics

Total number of embryos and TCs quantified (n) are provided in the figure legends. Measurement were imported and treated in Microsoft Excel, where graphics were generated. Error bars graphics and ± in text denote Standard Error of the Mean (SEM) or Standard deviation (SD). Statistical analysis were performed applying the T-test. Differences were considered significant when p<0,05. In graphics; *p<0.05, **p<0.01, ***p< 0.001.

### Time-lapse imaging

Dechorionated embryos were immobilised with glue on a coverslip and covered with Oil 10-S Voltalef (VWR). To visualise tracheal Shot *in vivo*, *btlGAL4UASShotC-GFP* was used in the indicated backgrounds. Actin in tracheal cells was visualised with *btl::moeRFP* or *btlGAL4UASlifeActRFP* where indicated. Imaging was done with a spectral confocal microscope Leica TCS SP5. The images were acquired for the times specified over 50-75 µm from st. 15 embryos; Z-projections and movies were assembled using Fiji (54).

## ACKNOWLEDGMENTS

We are grateful to M. Llimargas and our lab colleagues for comments on the manuscript. We thank J. Casanova and F. Serras for the support given during throughout this study. We thank A. Prokop, N. Sanchez-Soriano, K. Roeper, F. Jankovics and the Bloomington Drosophila Stock Center (BDSC) for fly stocks and reagents. Thanks also go to L. Bardia, A. Lladó, N. Giakoumakis and J. Colombelli from the IRB-ADMF for assistance and advice with confocal microscopy and software; C. Stephan-Otto Attolini from the IRB Bioinformatics/Biostatistics Facility; E. Fuentes, R. Mendez and M. Lledós for assistance in some experiments. D.R. is the recipient of a Juan de la Cierva post-doctoral fellowship from the Spanish *Ministerio de Ciencia, Innovación y Universidades* (FJCI201732443) and was previously funded by and FPU fellowship (FPU12/05765). This work was supported by the Universitat de Barcelona, Generalitat de Catalunya (2017 SGR 1455) and the Spanish *Ministerio de Ciencia, Innovación y Universidades* (PGC2018-099465-B-I00).

## Author contributions

Conceptualization: SA; Methodology and Experiments: DR and SA; Analysis: DR and SA; Writing, review and editing: DR and SA; Supervision and Funding: SA.

## REFERENCES

1. Gervais L, Casanova J. In vivo coupling of cell elongation and lumen formation in a single cell. Curr Biol. 2010;20(4):359–66.

2. Sigurbjörnsdóttir S, Mathew R, Leptin M. Molecular mechanisms of de novo lumen formation. Nat Rev Mol Cell Biol. 2014;15(10):665–76.

3. Best BT. Single-cell branching morphogenesis in the Drosophila trachea. Developmental Biology. 2019;451(1):5–15.

4. Affolter M, Montagne J, Walldorf U, Groppe J, Kloter U, LaRosa M, et al. The Drosophila SRF homolog is expressed in a subset of tracheal cells and maps within a genomic region required for tracheal development. Development. 1994;120(4):743–53.

5. Posern G, Treisman R. Actin’ together: serum response factor, its cofactors and the link to signal transduction. Trends in Cell Biology. 2006;16(11):588–96.

6. Lebreton G, Casanova J. Specification of leading and trailing cell features during collective migration in the Drosophila trachea. Journal of Cell Science. 2014;127(Pt 2):465–74.

7. Fischer RS, Lam P-Y, Huttenlocher A, Waterman CM. Filopodia and focal adhesions: An integrated system driving branching morphogenesis in neuronal pathfinding and angiogenesis. Developmental Biology. 2019;451(1):86–95.

8. Ricolo D, Deligiannaki M, Casanova J, Araújo SJ. Centrosome Amplification Increases Single-Cell Branching in Post-mitotic Cells. Curr Biol. 2016;26(20):2805–13.

9. Guillemin K, Groppe J, Ducker K, Treisman R, Hafen E, Affolter M, et al. The pruned gene encodes the Drosophila serum response factor and regulates cytoplasmic outgrowth during terminal branching of the tracheal system. Development. 1996;122(5):1353–62.

10. Schottenfeld-Roames J, Rosa JB, Ghabrial AS. Seamless Tube Shape Is Constrained by Endocytosis-Dependent Regulation of Active Moesin. Curr Biol. 2014;24(15):1756–64.

11. Oshima K, Takeda M, Kuranaga E, Ueda R, Aigaki T, Miura M, et al. IKK epsilon regulates F actin assembly and interacts with Drosophila IAP1 in cellular morphogenesis. Curr Biol. 2006;16(15):1531–7.

12. Okenve-Ramos P, Llimargas M. Fascin links Btl/FGFR signalling to the actin cytoskeleton during Drosophila tracheal morphogenesis. Development. 2014;141(4):929–39.

13. Jayanandanan N, Mathew R, Leptin M. Guidance of subcellular tubulogenesis by actin under the control of a synaptotagmin-like protein and Moesin. Nature Communications. 2014;5:3036.

14. Whitten JM. The Post-embryonic Development of the Trachea! System in Drosophila melanogaster. Journal of Cell Science. 1957;s3-98(41):1–30.

15. Baer MM, Bilstein A, Leptin M. A clonal genetic screen for mutants causing defects in larval tracheal morphogenesis in Drosophila. Genetics. 2007;176(4):2279.

16. Ghabrial AS, Levi BP, Krasnow MA. A Systematic Screen for Tube Morphogenesis and Branching Genes in the Drosophila Tracheal System. PLoS Genet. 2011;7(7):e1002087.

17. Jarecki J, Johnson E, Krasnow MA. Oxygen regulation of airway branching in Drosophila is mediated by branchless FGF. Cell. 1999;99(2):211–20.

18. Dogterom M, Koenderink GH. Actin-microtubule crosstalk in cell biology. Nature Reviews Molecular Cell Biology. 2019;20(1):38–54.

19. Suozzi KC, Wu X, Fuchs E. Spectraplakins: master orchestrators of cytoskeletal dynamics. The Journal of Cell Biology. 2012;197(4):465–75.

20. Röper K, Gregory SL, Brown NH. The ‘spectraplakins’: cytoskeletal giants with characteristics of both spectrin and plakin families. Journal of Cell Science. 2002;115(Pt 22):4215–25.

21. Gregory SL, Brown NH. kakapo, a gene required for adhesion between and within cell layers in Drosophila, encodes a large cytoskeletal linker protein related to plectin and dystrophin. The Journal of Cell Biology. 1998;143(5):1271–82.

22. Lee S, Harris KL, Whitington PM, Kolodziej PA. short stop is allelic to kakapo, and encodes rod-like cytoskeletal-associated proteins required for axon extension. Journal of Neuroscience. 2000;20(3):1096–108.

23. Applewhite DA, Grode KD, Keller D, Zadeh AD, Zadeh A, Slep KC, et al. The spectraplakin Short stop is an actin-microtubule cross-linker that contributes to organization of the microtubule network. Mol Biol Cell. 2010;21(10):1714–24.

24. Lee S, Kolodziej PA. Short Stop provides an essential link between F-actin and microtubules during axon extension. Development. 2002;129(5):1195–204.

25. Lee S, Kolodziej PA. The plakin Short Stop and the RhoA GTPase are required for E-cadherin-dependent apical surface remodeling during tracheal tube fusion. Development. 2002;129(6):1509–20.

26. Khanal I, Elbediwy A, Diaz de la Loza MdC, Fletcher GC, Thompson BJ. Shot and Patronin polarise microtubules to direct membrane traffic and biogenesis of microvilli in epithelia. Journal of Cell Science. 2016;129(13):2651–9.

27. Subramanian A, Prokop A, Yamamoto M, Sugimura K, Uemura T, Betschinger J, et al. Shortstop recruits EB1/APC1 and promotes microtubule assembly at the muscle-tendon junction. Curr Biol. 2003;13(13):1086–95.

28. Sun T, Song Y, Dai J, Mao D, Ma M, Ni J-Q, et al. Spectraplakin Shot Maintains Perinuclear Microtubule Organization in Drosophila Polyploid Cells. Developmental Cell. 2019.

29. Mui UN, Lubczyk CM, Nam S-C. Role of spectraplakin in Drosophila photoreceptor morphogenesis. PLoS ONE. 2011;6(10):e25965.

30. Nashchekin D, Fernandes AR, St Johnston D. Patronin/Shot Cortical Foci Assemble the Noncentrosomal Microtubule Array that Specifies the Drosophila Anterior-Posterior Axis. Developmental Cell. 2016;38(1):61–72.

31. Röper K, Brown NH. Maintaining epithelial integrity: a function for gigantic spectraplakin isoforms in adherens junctions. The Journal of Cell Biology. 2003;162(7):1305–15.

32. Gervais L, Casanova J. The Drosophila homologue of SRF acts as a boosting mechanism to sustain FGF-induced terminal branching in the tracheal system. Development. 2011;138(7):1269–74.

33. Bottenberg W, Sánchez-Soriano N, Alves-Silva J, Hahn I, Mende M, Prokop A. Context-specific requirements of functional domains of the Spectraplakin Short stop in vivo. Mechanisms of Development. 2009;126(7):489–502.

34. Voelzmann A, Liew Y-T, Qu Y, Hahn I, Melero C, Sánchez-Soriano N, et al. Drosophila Short stop as a paradigm for the role and regulation of spectraplakins. Seminars in Cell and Developmental Biology. 2017:1–18.

35. Alves-Silva J, Sánchez-Soriano N, Beaven R, Klein M, Parkin J, Millard TH, et al. Spectraplakins promote microtubule-mediated axonal growth by functioning as structural microtubule-associated proteins and EB1-dependent +TIPs (tip interacting proteins). Journal of Neuroscience. 2012;32(27):9143–58.

36. Hahn I, Ronshaugen M, Sánchez-Soriano N, Prokop A. Functional and Genetic Analysis of Spectraplakins in Drosophila. Meth Enzymol. 2016;569:373–405.

37. Sanchez-Soriano N, Travis M, Dajas-Bailador F, Goncalves-Pimentel C, Whitmarsh AJ, Prokop A. Mouse ACF7 and Drosophila Short stop modulate filopodia formation and microtubule organisation during neuronal growth. Journal of Cell Science. 2009;122(14):2534–42.

38. Takács Z, Jankovics F, Vilmos P, Lénárt P, Röper K, Erdélyi M. The spectraplakin Short stop is an essential microtubule regulator involved in epithelial closure in Drosophila. Journal of Cell Science. 2017;130(4):712–24.

39. Olson EN, Nordheim A. Linking actin dynamics and gene transcription to drive cellular motile functions. Nat Rev Mol Cell Biol. 2010;11(5):353–65.

40. Blanco E, Messeguer X, Smith TF, Guigo R. Transcription factor map alignment of promoter regions. PLoS Comput Biol. 2006;2(5):e49.

41. Khan A, Fornes O, Arnaud Stigliani MG, Jaime A Castro-Mondragon, Robin van der Lee, Adrien Bessy, Jeanne Chèneby, Shubhada R Kulkarni, Ge Tan, Damir Baranasic, David J Arenillas, Albin Sandelin, Klaas Vandepoele, Boris Lenhard, Benoît Ballester, Wyeth W Wasserman, François Parcy, Anthony Mathelier JASPAR 2018: update of the open-access database of transcription factor binding profiles and its web framework. Nucleic Acids Res. 2018;46:D260–D6.

42. Voelzmann A, Okenve-Ramos P, Qu Y, Chojnowska-Monga M, Del Caño-Espinel M, Prokop A, et al. Tau and spectraplakins promote synapse formation and maintenance through Jun kinase and neuronal trafficking. eLife. 2016;5:2322.

43. Prout M, Damania Z, Soong J, Fristrom D, Fristrom JW. Autosomal mutations affecting adhesion between wing surfaces in Drosophila melanogaster. Genetics. 1997;146(1):275–85.

44. Lee M, Nahm M, Kwon M, Kim E, Zadeh AD, Cao H, et al. The F-actin-microtubule crosslinker Shot is a platform for Krasavietz-mediated translational regulation of midline axon repulsion. Development. 2007;134(9):1767–77.

45. Sutherland D, Samakovlis C, Krasnow MA. branchless encodes a Drosophila FGF homolog that controls tracheal cell migration and the pattern of branching. Cell. 1996;87(6):1091–101.

46. Riederer BM. Microtubule-associated protein 1B, a growth-associated and phosphorylated scaffold protein. Brain Research Bulletin. 2007;71(6):541–58.

47. Morris M, Maeda S, Vossel K, Mucke L. The many faces of tau. Neuron. 2011;70(3):410–26.

48. Elie A, Prezel E, Guérin C, Denarier E, Ramirez-Rios S, Serre L, et al. Tau co-organizes dynamic microtubule and actin networks. Scientific Reports. 2015;5:9964.

49. Cabrales Fontela Y, Kadavath H, Biernat J, Riedel D, Mandelkow E, Zweckstetter M. Multivalent cross-linking of actin filaments and microtubules through the microtubule-associated protein Tau. Nature Communications. 2017;8(1):1981–12.

50. Biswas S, Kalil K. The Microtubule-Associated Protein Tau Mediates the Organization of Microtubules and Their Dynamic Exploration of Actin-Rich Lamellipodia and Filopodia of Cortical Growth Cones. Journal of Neuroscience. 2018;38(2):291–307.

51. Lee T, Hacohen N, Krasnow M, Montell DJ. Regulated Breathless receptor tyrosine kinase activity required to pattern cell migration and branching in the Drosophila tracheal system. Genes Dev. 1996;10(22):2912–21.

52. Levi BP, Ghabrial AS, Krasnow MA. Drosophila talin and integrin genes are required for maintenance of tracheal terminal branches and luminal organization. Development. 2006;133(12):2383–93.

53. Zhang J, Yue J, Wu X. Spectraplakin family proteins - cytoskeletal crosslinkers with versatile roles. Journal of Cell Science. 2017;130(15):2447–57.

54. Schindelin J A-CI, Frise E, Kaynig V, Longair M, Pietzsch T, Preibisch S, Rueden C, Saalfeld S, Schmid B, Tinevez JY, White DJ, Hartenstein V, Eliceiri K, Tomancak P, Cardona A. Fiji: an open-source platform for biological-image analysis. Nat Methods. 2012;9(7):676–82.

