## Supplementary Information for "Coordinated crosstalk between microtubules and actin by a spectraplakin regulates lumen formation and branching"

### SUPPLEMENTARY MATERIALS

**Supplementary Table 1.** DSRF-specific sequences in upstream and downstream regions of the *shot* gene. Sequences are shown within 2000 bp of the TSS with more than 70% similarity to the specific DSRF binding sequences, indicating their score and the chain of chromosome 2 in which they were found.

The DSRF-binding sequences identified, overlap with regions P1, P2, and P3. In the case of the P1 promoter, we found the sequences in the table spanning 13,926,470-13,927,188; in the case of P2, the first three DSRF binding sequences of the table are located within this region, which include 13,943,065-13,329; and in the case of P3, we found DSRF binding sequences within 13,912,773-13,912,789.

| Chr | Start | End | Length | Score | Chain | Sequence |
|---|---|---|---|---|---|---|
| chr2R | 13,943,313 | 13,943,329 | 16 | 0.75 | + | TTTCCAAATGGGGTCA |
| chr2R | 13,943,313 | 13,943,329 | 16 | 0.75 | - | TGACCCCATTTGGAAA |
| chr2R | 13,943,065 | 13,943,081 | 16 | 0.74 | + | TCCCGATCTGTGGTCA |
| chr2R | 13,927,172 | 13,927,188 | 16 | 0.78 | + | TACTCATTTATGGACA |
| chr2R | 13,927,172 | 13,927,188 | 16 | 0.76 | - | TGTCCATAAATGAGTA |
| chr2R | 13,926,681 | 13,926,697 | 16 | 0.74 | + | TGTCGATCTGTGGTCT |
| chr2R | 13,926,470 | 13,926,486 | 16 | 0.79 | + | TGACCAAATATGAGGA |
| chr2R | 13,926,470 | 13,926,486 | 16 | 0.80 | - | TCCTCATATTTGGTCA |
| chr2R | 13,912,773 | 13,912,789 | 16 | 0.76 | + | ACTCCATATAGGGCAA |
| chr2R | 13,912,260 | 13,912,276 | 16 | 0.73 | + | TTAACATAAGTGGTCA |
| chr2R | 13,912,260 | 13,912,276 | 16 | 0.74 | - | TGACCACTTATGTTAA |
| chr2R | 13,912,773 | 13,912,789 | 16 | 0.78 | - | TTGCCCTATATGGAGT |

**Supplementary Table 2. DSRF positional weight matrix (PWM)**

Specific DSRF-binding sequences known in vertebrates (Khan et al., 2018).

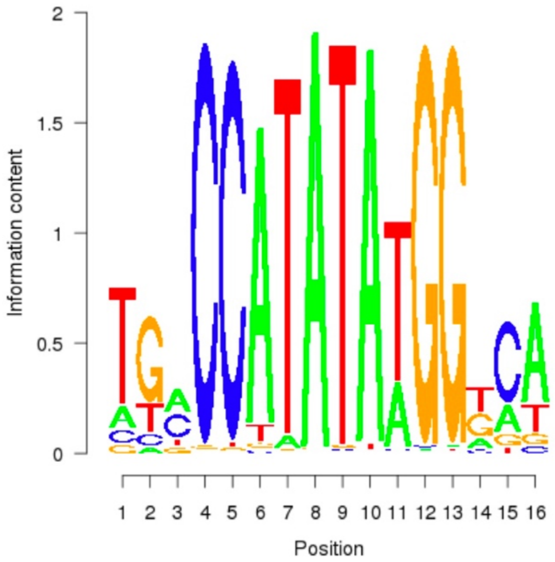


**Movie 1 – Shot localization within the TCs is dynamic and accompanies the growing subcellular lumen.** *In vivo* Shot localization during lumen formation in two wild-type ganglionic branch (GB) TCs. Time-lapse images of a *wt* embryo expressing btlGAL4UASShotC-GFP visualized from a dorsal view. Note the accumulation of Shot at the apical junction of the TC and subsequently in association with subcellular lumen extension. Frames were taken every minute for 3,5 hours.

**Movie 2 - Shot localizes with moesin in TCs during subcellular lumen extension.** *In vivo* Shot colocalization with actin during lumen formation in a wild-type dorsal branch (DB) TC. Time-lapse images of a *wt* embryo expressing btlGAL4UASShotC-GFP; btl::moeRFP visualized from a dorsal view. Note the colocalization of Shot with actin the apical junction of the TC and subsequently in association with the actin core and filopodial actin during subcellular lumen extension. Frames were taken every 2 minutes for 3,3 hours.

**Movie 3 - Shot localizes with actin in TCs during subcellular lumen extension.** *In vivo* Shot colocalization with actin during lumen formation in a wild-type dorsal branch (DB) TC. Time-lapse images of a *wt* embryo expressing btlGAL4UASShot-C-GFPUASlifeActRFP visualized from a dorsal view. Note the colocalization of Shot with actin the apical junction of the TC and subsequently in association with the actin core and filopodial actin during subcellular lumen extension. Frames were taken every 40 seconds for 32 minutes.

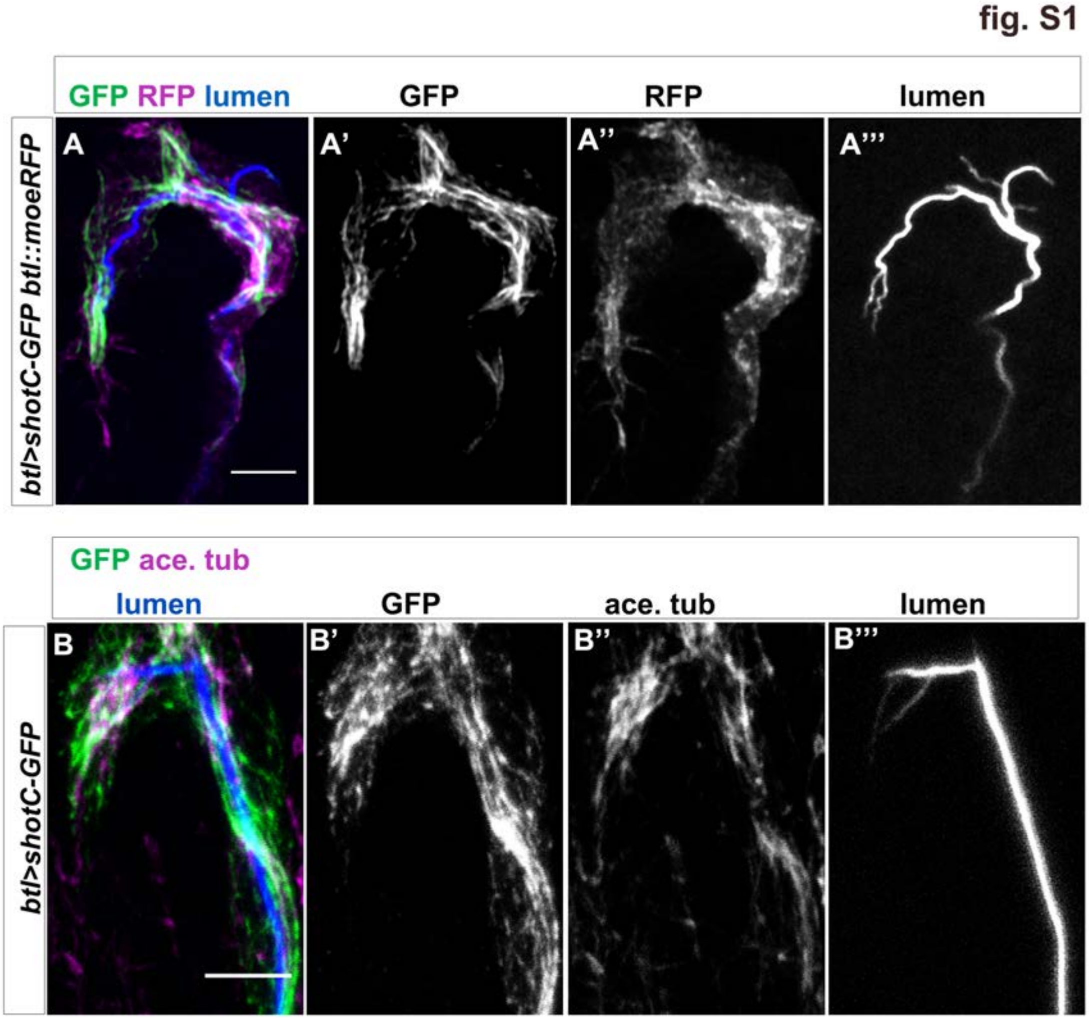


**Figure S1. ShotOE does not perturb the TC cytoskeletal organization**.

(A) Dorsal TC of a late st.15 *btl>shotCGFP; btl::moeRFP* embryo stained with GFP (green in A, grey in A'), RFP (magenta in A grey in A'' and CBP (blue in A, grey in A'''). The extra subcellular lumen appears inside the same cytoplasmatic protrusion in 75% of TCs analysed (n=25).

(B) Dorsal TCs of an early st. 15 *btl>ShotC-GFP* stained with GFP (green in B, grey in B'), acetylated-tubulin (magenta in B, grey in B'') and CBP (blue in B grey in B'''). GFP positive bundles localize with MTs (n=20). Scale bar 5µm.

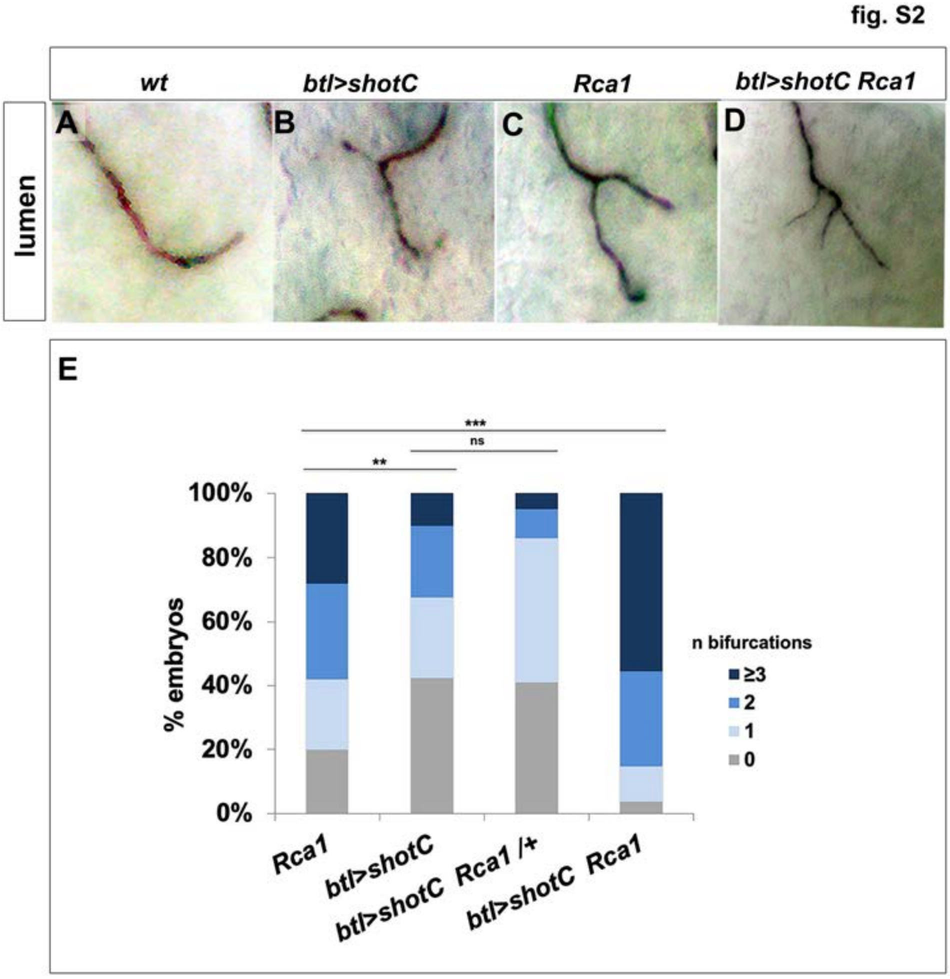


**Figure S2. ESL induction by *ShotOE* is additive to centrosome amplification.**

(A-D) Details of GB TCs at st. 16 from embryos stained with anti-Gasp antibody to mark the lumen. (A) *wt* TCs with a single lumen each; (B) *btl>shotC* showing subcellular lumen bifurcations; (C) *Rca1* showing subcellular lumen bifurcation; (D) *Rca1; btl>shotC* showing a multi-branched subcellular lumen. Anterior side of the embryo is on the left ventral midline is down.

(E) Quantification of the number of bifurcations (GB TCs) per embryo of the indicated genotype. In the graph embryos with 0 bifurcations are represented by grey columns, embryos with 1 bifurcation columns in light blue, 2 bifurcations columns in blue, 3/more bifurcations columns in dark blue. To better quantify the phenotypes, we subdivided *Rca1* and ShotOE embryos in groups characterized by 1, 2, “3/or more” branched TC lumina. 80% of *Rca1* mutant embryos had TCs with supernumerary lumina (n=40); in particular, 24% of embryos had 1 bifurcation, 30% of embryos had 2 bifurcations and 28% of embryos had 3 or more bifurcations. In ShotOE embryos, 57,5 % of embryos had TCs affected; 25% with one bifurcation, 22.5% with two bifurcations and 10% with three or more bifurcations (n= 40). We then analysed the phenotype in *Rca1,* ShotOE embryos (n=28). These embryos had bifurcation phenotypes in 96,3 % of the cases; 11,1% with 1 bifurcation 29,6% with 2 bifurcations and 55,6% with 3 or more bifurcations. So, in *Rca1,* ShotOE embryos, we observed a higher number of embryos with 2 or 3/more bifurcations and a lower number of embryos with 0 or 1 bifurcations in relation to *Rca1*, suggesting that the effect of *Rca1* LOF and ShotOE was additive in producing 2 or 3/more bifurcations.

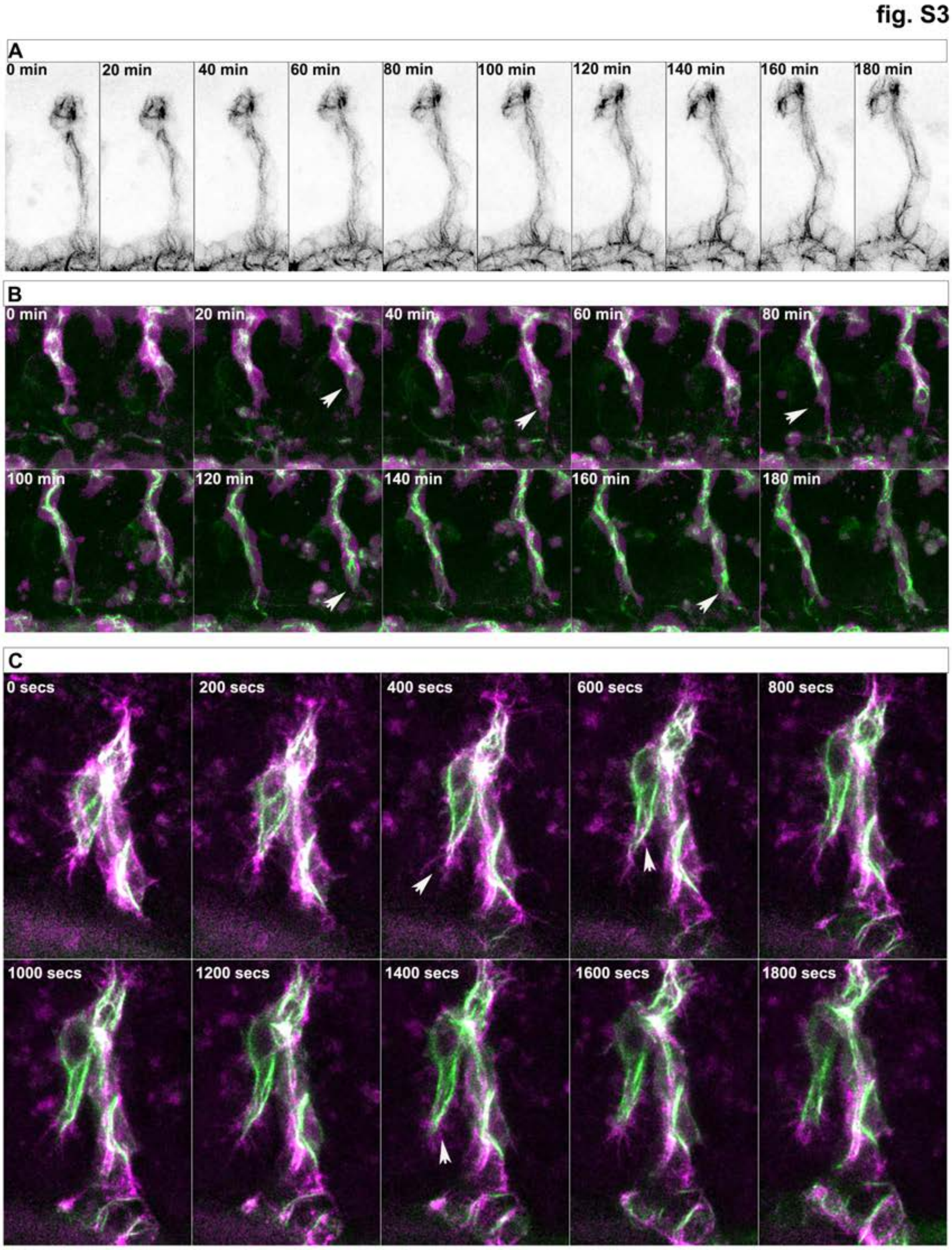


**Figure S3 – Shot is dynamically localized in TCs and interacts with Moe and actin.** (A) Movie 1 frames from the beginning of image acquisition to 180 min. Live embryos expressing ShotC-GFP in all tracheal cells. (B) Movie 2 frames from the beginning of image acquisition to 180 min. Live embryos expressing ShotC-GFP and MoeRFP in all tracheal cells. (C) Movie 3 frames from the beginning of image acquisition to 30 min. Live embryos expressing ShotC-GFP and lifeActinRFP in all tracheal cells.

**fig.S4**

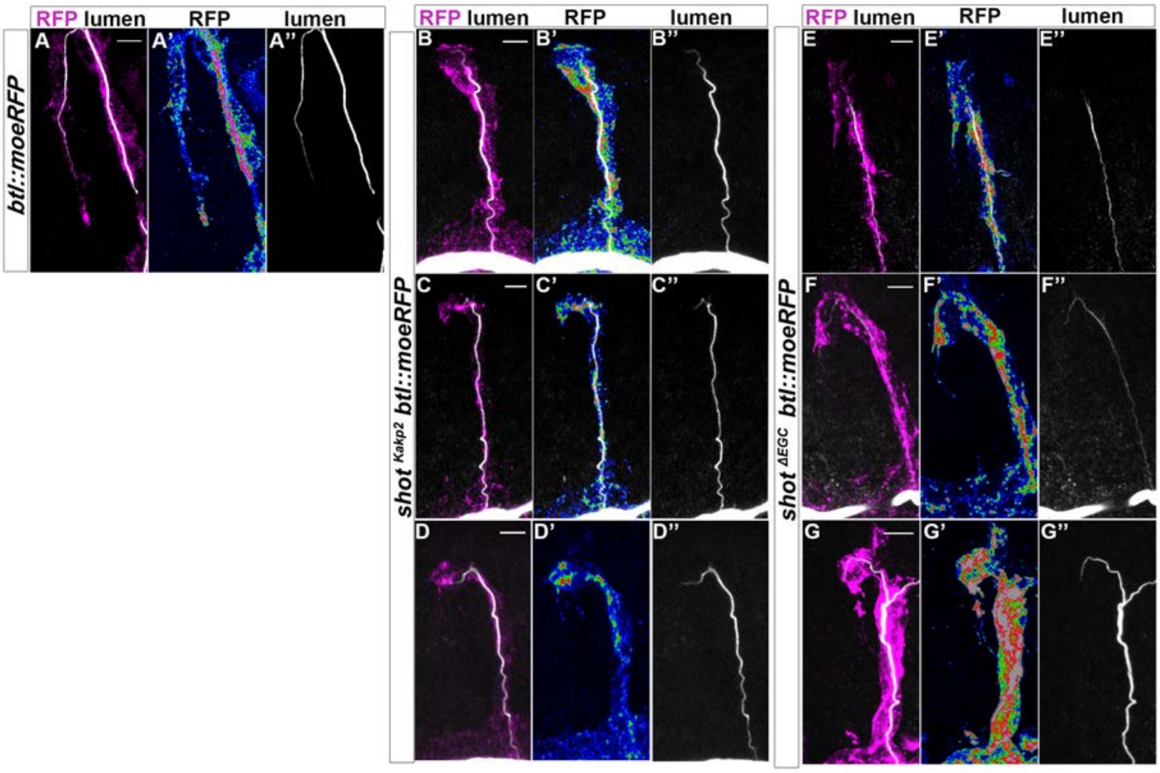


**Figure S4. Asymmetric actin accumulation is affected in *shot* $^{kakp2}$ and *shot* $^{\Delta EGC}$.**

Dorsal TCs from *btl::moeRFP* (A) *shot*$^{KAKP2}$ *; btl::moeRFP* (B-D) and *shot*$^{\Delta EGC}$ *; btl::moeRFP* (E-G) embryos stained with RFP (magenta in A-G, in a colour scale in which blue is low, green in middle and red high pixel intensity (A'-G') and CBP in grey (A-G and A''-G''). Similarly to the null allele *shot*$^{3}$*,* actin organization was impaired in all cases in both mutants (ranging from partially elongated TCs with subcellular lumen to cases in which the cell did not elongate and a subcellular lumen was not formed). Scale bar 5 µm.

fig. S5

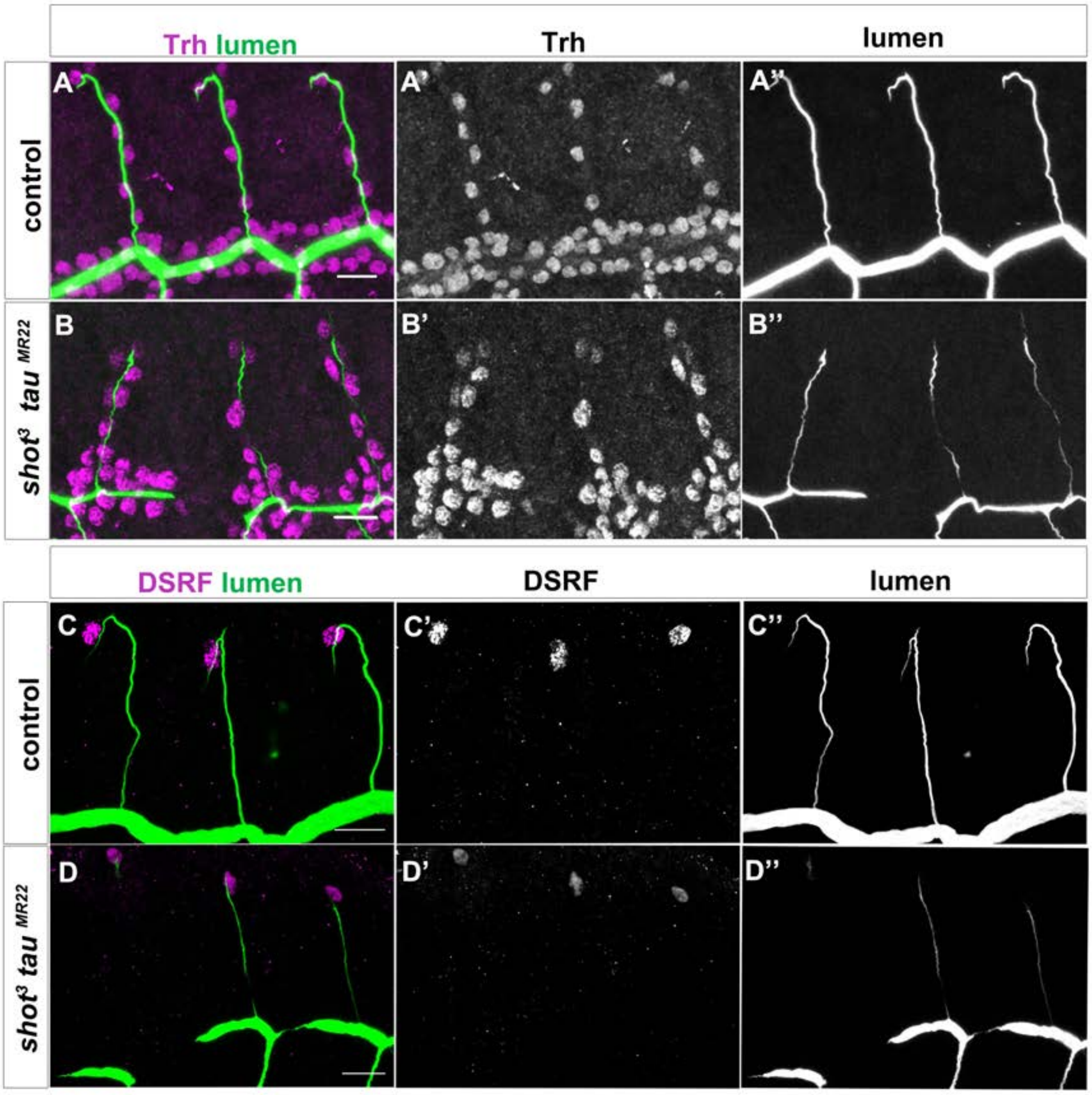


**Figure S5. Double *shot³;tau^MR22^* mutant embryos display defects in lumen formation but do not in tracheal cell number or TC fate**.

Dorsal branches of *wt* (A) and *shot³; tau [MR22]* embryos st.15 (B) stained with trachealess (trh) antibody to visualize all tracheal cell nuclei (magenta in A and B, grey in A' and B') and CBP to visualize the lumen (green in A and B, grey in A'' and B'' (n= 17 TCs) or wt (C) and *shot³; tau [MR22]* (D) stained with DSRF magenta in C and D (grey in C' and D') and CBP green in C and D and grey in C'' and D'' (n= 8 TCs). Dorsal view, scale bar 10 μm.

Khan, A., O. Fornes, and M.G. Arnaud Stigliani, Jaime A Castro-Mondragon, Robin van der Lee, Adrien Bessy, Jeanne Chèneby, Shubhada R Kulkarni, Ge Tan, Damir Baranasic, David J Arenillas, Albin Sandelin, Klaas Vandepoele, Boris Lenhard, Benoît Ballester, Wyeth W Wasserman, François Parcy, Anthony Mathelier 2018. JASPAR 2018: update of the open-access database of transcription factor binding profiles and its web framework. *Nucleic Acids Res.* 46:D260-D266.
